# Sex-specific differences in endocannabinoid regulation of cocaine-evoked dopamine in the medial nucleus accumbens shell

**DOI:** 10.64898/2026.03.27.714857

**Authors:** Andrew D. Gaulden, Kristin Chase, Jayme R. McReynolds

## Abstract

Endocannabinoid (eCB) signaling is a key regulator of reward-related dopaminergic signaling, particularly in response to drugs of abuse, such as cocaine. To date, our understanding of this mechanism has primarily been limited to male subjects. Prior work establishes that female cocaine users have more adverse outcomes, and female rats show greater sensitivity to cannabinoid type 1 receptor (CB1R) regulation of cocaine self-administration. Therefore, we hypothesize that female rats exhibit enhanced eCB regulation of cocaine-evoked dopamine (DA). We used *in vivo* fiber photometry recording of the dopamine biosensor, dLight 1.3b, in the nucleus accumbens medial shell (NAc_ms_) in response to cocaine in male and female rats. Rats were pretreated with cannabinoid-targeting drugs to investigate the effects of CB1R inactivation or augmentation of the eCB 2-AG on cocaine-evoked DA. Our results revealed that CB1R inactivation attenuates cocaine-evoked DA in male and female rats, but females showed enhanced sensitivity for CB1R regulation of cocaine-evoked DA. Cocaine-evoked DA was enhanced by augmenting 2-AG levels, and females again showed increased sensitivity to this manipulation. Finally, females show greater cocaine-evoked DA when in a non-estrous cycle compared to estrous, reinforcing that estrous cycle is a determinant of cocaine-evoked DA. These data indicate that females show enhanced eCB regulation of cocaine-evoked DA signaling, underscoring the importance of sex as a biological variable in our understanding of endocannabinoid regulation of drug reward.

**Highlights:**

- CB1R inactivation attenuates cocaine-evoked DA in NAc_ms_, preferentially in females
- 2-AG augmentation via MAGL inhibition enhances cocaine-evoked DA, with female bias
- Estrous phase modulates the dopamine response to a high dose of cocaine in females
- Male and female rats show similar baseline DA and locomotor responses to cocaine

## 1. Introduction

Cocaine use disorder (CUD) remains a critical issue in the United States with more than 1.3 million Americans having been recently diagnosed with CUD (*Substance Abuse and Mental Health Services Administration*, 2022). Despite years of research, there is still no FDA-approved medication for the treatment of CUD, making this a critical unmet need. Therefore, to develop better therapeutic strategies it is imperative to fully understand how the reinforcing efficacy of cocaine is regulated. Cocaine exerts its primary reinforcing efficacy through elevation of extracellular dopamine (DA) in the nucleus accumbens (NAc; Jones et al., 1995; Woolverton & Johnson, 1992). However, pharmacotherapies for individuals with CUD that target DA signaling have largely failed in clinical trials (Brandt et al., 2021; Chan et al., 2019). Therefore, it is important to identify other signaling systems that can regulate the dopaminergic response to cocaine in reward-related brain regions, either through regulation of upstream ventral tegmental area (VTA) DA neuron firing or regulation of downstream extracellular DA levels in the NAc.

One system of recent interest in the regulation of drugs of abuse is the endocannabinoid (eCB) system (Solinas et al., 2008; Wenzel & Cheer, 2018). Endocannabinoids are retrograde neurotransmitters produced in the postsynaptic neuron that bind to a pre-synaptic G_i/o_-coupled cannabinoid type 1 receptor (CB1R) to inhibit neurotransmission (Kano et al., 2009). CB1Rs are localized on glutamatergic and GABAergic terminals in the VTA and NAc and tightly regulate DA signaling to modulate reward (Covey & Yocky, 2021; Parsons & Hurd, 2015; Riegel & Lupica, 2004). Numerous studies have identified that cocaine can mobilize 2-arachidonoylglycerol (2-AG), the most abundant eCB, in the VTA and disinhibit VTA DA neurons through CB1R activation on inhibitory terminals (Engi et al., 2021; Nakamura et al., 2019; Wang et al., 2015). Additionally, there is evidence that eCBs can regulate cocaine responses in the NAc shell either through regulation of glutamatergic inputs (Ingebretson et al., 2018) or fast spiking interneurons (Winters et al., 2012). Together, this work by Cheer and others provides a strong model for how cocaine-evoked DA signaling is facilitated by CB1R signaling (Zlebnik & Cheer, 2016). However, the CB1R enhancement of cocaine-evoked DA seen in these neurocircuitry investigations is not consistently supported in rodent models of cocaine-motivated behavior.

Overall, there is limited and inconsistent information supporting the hypothesis that CB1R signaling regulates cocaine-motivated behavior (Cossu et al., 2001; De Vries et al., 2001; Fattore et al., 2000; Vlachou et al., 2003; Wiskerke et al., 2008). There is evidence that CB1Rs may play more of a role in regulation of cocaine self-administration under conditions leading to dysregulated cocaine-motivated behavior (McReynolds et al., 2023; Orio et al., 2009). However, because CB1Rs are found throughout the brain, these disparate effects may also be due to whether eCB manipulation was done systemically or in a region-specific manner.

For example, in the VTA, cocaine self-administration upregulates CB1R binding and intra-VTA CB1R antagonism attenuates cocaine self-administration even under limited access conditions (McReynolds et al., 2023). Therefore, more work needs to be done to deeply interrogate how endocannabinoid signaling regulates the dopaminergic response to cocaine in mesolimbic brain regions.

Of the preclinical models investigating the contribution of CB1R signaling to cocaine reward, very few have directly investigated sex as a biological variable. Clinical literature indicates that women who use cocaine have more adverse outcomes than men (Becker & Chartoff, 2018; Becker & Hu, 2008), including elevated cocaine use (Schottenfeld et al., 1993). Sex differences in cocaine reward have been broadly attributed to gonadal hormones (Bobzean et al., 2014), and there is compelling evidence that estradiol and progesterone can influence cocaine reward in females (Calipari et al., 2017; Peart et al., 2022; Peris et al., 1991; Sofuoglu et al., 2004). Although estradiol has been shown to enhance cocaine reward through various mechanisms (Peart et al., 2022), there is evidence that estradiol utilizes CB1R signaling to promote a sensitized response to cocaine (Peterson et al., 2016) and facilitate cocaine self-administration (Martinez et al., 2016). Despite the evidence of CB1R signaling and sex as a biological variable interacting to promote cocaine reward, very little work has been done to elucidate how these factors affect reward circuitry. Our own work (Gaulden et al., 2025) indicates that females may show greater overall sensitivity to CB1R inactivation in the context of cocaine reward.

However, it is unclear to what extent eCB signaling regulates cocaine-evoked dopamine in the NAc medial shell in females, representing a critical gap in the field. Therefore, here we investigate the sex-specific effects of eCB regulation on cocaine-evoked dopamine in the nucleus accumbens shell using *in vivo* fiber photometry recording of a dopamine biosensor. This technique allows performing longitudinal recordings for the examination of eCB regulation of cocaine-evoked dopamine in a dose- and sex-dependent manner.

## 2. Methods

### 2.1. Subjects

Male (n = 24) and freely cycling female (n = 24) Long-Evans rats (approx. 70 days old; Males: 250-275 g at arrival; Females: 225-250g at arrival; Envigo, Indianapolis, IN) were housed individually in a temperature-and humidity-controlled AAALAC-accredited animal facility on a regular 12:12 hr light cycle (8 am lights on).

Rats were given *ad libitum* access to food and water. Procedures were approved by the Institutional Animal Care and Use Committee at the University of Cincinnati and conducted in compliance with NIH guidelines. All experiments were conducted during the light phase, approximately between 11:00 AM and 6:00 PM.

### 2.2. Drugs and Reagents

Cocaine HCl and Rimonabant were obtained from the National Institute of Drug Abuse (NIDA) Drug Supply Program. MJN-110 was acquired from MedChemExpress (Monmouth Junction, NJ). Cocaine was dissolved in bacteriostatic 0.9% saline and diluted to working stocks with concentrations of 0.5, 1, and 3 mg/mL for infusions. Raclopride was acquired from Tocris (Minneapolis, MN) and was dissolved in bacteriostatic 0.9% saline at a concentration of 2 mg/kg, (i.p). Propofol injectable emulsion (10 mg/ml) was sourced from Covetrus (Dublin, OH) and delivered at 0.2 ml/kg, i.v.. Doses for Rimonabant were selected based on our prior work and the work of others showing efficacy at reducing cocaine-taking behavior at 1 mg/kg, i.p. and 3 mg/kg, (i.p.) (Gaulden et al., 2025; Engi et al., 2021). Rimonabant was first dissolved in ethanol, followed by Kolliphor EL (Millipore Sigma, Burlington, MA) and finally by bacteriostatic saline (0.9%) in a 1:1:18 ratio. The vehicle consisted of the same 1:1:18 ratio of ethanol: Kolliphor EL: saline. Rimonabant pretreatment time was set at 25 minutes to allow for maximal effect of CB1R inactivation during photometry recording (Cheer et al., 2007; Gaulden et al., 2025; Rinaldi-Carmona et al., 1994). Dose for MJN-110 was selected based on prior work by Cravatt and colleagues (Niphakis et al., 2013; Wilkerson et al., 2016) which saw a maximal MAGL inhibition at 5 mg/kg, i.p. MJN-110 was prepared similarly to Rimonabant, but at a 1:1:8 ratio of ethanol: Kolliphor EL: saline. MJN pretreatment time was set at 175 minutes to allow for full MAGL inhibition and the gradual accumulation of 2-AG (Ignatowska-Jankowska et al., 2015).

### 2.3. Viral infusion and fiber optic cannula implantation

All rats received craniotomies for *in vivo* fiber photometry surgery. Rats were anesthetized with isoflurane (2-2.5% in O2; Covetrus North America, Portland, ME) and mounted into a stereotaxic frame (Kopf, Tujunga, CA). The skull surface was cleared of any connective tissue, and four skull screws (McMaster Carr,

Elmhurst, IL) were anchored to the skull around the surgical site. A unilateral craniotomy was performed over the NAc_ms_ (From Bregma: Anterior/Posterior (A/P): + 1.80 mm, Medial/Lateral (M/L): ±0.95 mm, Dorsal/Ventral (D/V): - 7.00 mm) to infuse dLight 1.3b into the NAc_ms_ (∼5.2 × 10^12^ vg/mL, AAV5-CAG-dLight1.3b was a gift from Lin Tian (Addgene viral prep # 111067-AAV5; http://n2t.net/addgene:111067 ; RRID:Addgene_111067)). A unilateral 500 nL microinjection of dLight 1.3b was infused at the NAc_ms_ coordinates, and another 500 nL was infused 0.2 mm above the site to ensure maximal spread. After each infusion, an 8 min wait time was allowed for the viral bolus to adequately diffuse. Next, a unilateral fiber optic cannula (400-µm diameter core, 0.48 numerical aperture, 2.5 mm metal ferrule; Doric Lenses, Quebec, Canada) was lowered to an offset position from the injection site to avoid tissue damage from the microinfusion (From Bregma: A/P: + 1.80 mm, M/L: ±1.10 mm, D/V: -6.80 mm). The fiber optic cannula was implanted to the skull surface first with a thin layer of C&B Metabond (Parkell, Inc, Edgewood, NY), then the implant was finished using acrylic dental cement (Stoelting, Wood Dale, IL). The side of the dLight 1.3b infusion and fiber optic cannula implantation was counterbalanced across rats. During anesthesia, and for 2 days post-surgery, rats received daily carprofen injections (5 mg/kg, s.c.; Covetrus, Dublin, OH). Three weeks after *in vivo* fiber photometry surgery, rats were tested for physiologically relevant photometric signal. Rats that showed a positive photometric signal were later implanted with an intravenous catheter for behavioral testing.

### 2.4. Intravenous catheter implantation

Rats were anesthetized with isoflurane (2-2.5% in O2; Covetrus North America, Portland, ME) and the catheter was implanted into the superior vena cava with the back mount situated approx. 2-2.5 cm behind the rat’s scapula. The catheter consisted of a back-mounted cannula (Plastics One, Roanoke, VA) connected on the underside of the back mount to polypropylene mesh (500 microns; Small Parts, Logansport, IN). The back mount cannula was attached to a polyurethane catheter (0.6 mm i.d. x 1.1 mm o.d.; Access Technologies, Skokie, Il). Rats were given carprofen (5 mg/kg, sc) at the onset of the surgery and for 2 additional days. The catheters were flushed with Cefazolin antibiotic treatment (100 mg/kg, i.p.) in heparinized bacteriostatic 0.9% saline for 6 days following surgery. Following surgery recovery, the Cefazolin treatment ceased but the catheters continued to be flushed daily with heparinized bacteriostatic 0.9% saline throughout the experiment. Rats recovered for a minimum of one week before the initiation of behavioral testing.

### 2.5. Fiber Photometry Recordings

To excite and record from the dLight 1.3b biosensor, we used a commercially available photometry hardware and software package (Tucker Davis Technologies, Alachua, FL). The RZ10X processor and amplifier, with LEDs emitting light at wavelengths for excitation of GFP and for isosbestic control (465 nm and 405 nm, respectively) delivered to two minicubes (Doric Lenses, Quebec, Canada), was used to transmit light and receive and record dLight 1.3b fluorescent signal. The collected signal was processed with Synapse software suite (Tucker Davis Technologies, Alachua, FL). For every recording day, LEDs were powered with sufficient mV to reach 30uW per the RZ10x integrated power meter. Prior to recordings, optical cables were photobleached overnight for 4h, twice per week. Time stamps for epochs during photometry recordings were made via experimenter-triggered software buttons in Synapse. To couple the fiber optic implant to the optical patch cable (Ø 400 µm, NA 0.37, stainless steel armor jacket; Doric Lenses, Quebec, Canada), 2.5 mm diameter ceramic mating sleeves (Thorlabs, Newton, NJ) were used and the connection was firmly tightened with a custom 3D-printed clamp and metal lug screw. Rats were video recorded during testing with overhead high-definition USB webcams that were integrated with Synapse to record at a fixed frame rate of 20 FPS.

### 2.6. Fiber Photometry Analysis

Photometry data were extracted and analyzed using two different Python-based graphical user interfaces (GUIs): GuPPY (Sherathiya et al., 2021) and custom Python code in Stremecoder (Pluricorp, Tustin, CA). In either case, the isosbestic signal from the 405 nm channel was filtered with a low-pass (10Hz) filter, the 465 nm signal was then de-trended using a polyfit line of the 405 nm signal, and finally the change in fluorescence of the biosensor at 465 nm was divided by the control 405 nm signal to yield ΔF (per Learner et al., 2015). The ΔF was then divided by the control channel value for each time point to yield ΔF/f. The processed data was downsampled to a rate of 100Hz for easier graphing and data management. To extract epochs from user inputs and time-lock them to the ΔF/f of fluorescent activity, we used Stremecoder, a Python-based customizable program. To analyze changes in baseline transient amplitude and frequency after drug injections, we used GuPPY (Sherathiya et al., 2021). The Z score threshold was set at 2, and any drops in signal reflected in the 405 nm isosbestic channel were excluded and the remaining signal was concatenated.

To mitigate individual differences in dynamic range of photometry signals, we conducted Z-score assessment of our photometry traces. To calculate Z-scored values, the standard deviation for Z = 0 was collected from a 30 s baseline of ΔF/f values prior to the infusion. This baseline did not include data from 15s prior to infusion to account for transient occurrences related to experimenter handling of the animal in preparation for infusion delivery. Area under the curve (AUC) was calculated for Z-normalized dLight signal for 90s immediately following the infusion. In most cases, AUC analysis was conducted for three 30s time bins during the 90s post-infusion period and in a small number of experiments AUC analysis was conducted for the entire 90s post-infusion period. We also analyzed the maximum peak of our down-sampled, Z-scored data. This metric captured the maximal Z-score within the 90s post-infusion period.

To assess potential order effects within the testing day, we examined the change in the dLight signal during the post-infusion period from the first infusion of cocaine to the second infusion of cocaine within the same testing day in each rat. Magnitude changes in response were based on AUC data of an initial dose of cocaine followed by the subsequent matched dose for the day. Mathematically, magnitude change was derived as: [(second infusion AUC / first infusion AUC) * 100] -100. We then compared the magnitude change (from first infusion to second infusion within the day) of repeated cocaine infusions alone or in the presence of cannabinoid modulator drugs. Final graphs were plotted with GraphPad Prism (GraphPad Software Inc., San Diego, CA).

### 2.7. Vaginal Lavage Cytology

We used estrous cycle staging to indirectly assess circulating gonadal hormones in our female rats. Based on prior publications, we hypothesized that estradiol regulates cocaine-evoked DA signaling in the NAc_ms_. To assess this relationship, we collected vaginal lavage samples on test days when the dLight response to cocaine was recorded. Using a P200 pipette, the vaginal canal was flushed with distilled water, and the cell-containing water was collected onto microscope slides and immediately stained with Cresyl Violet (MilliporeSigma, Burlington, MA). The vaginal lavage samples were analyzed for estrous cycle phases using published methods (McLean et al., 2012; Cora et al., 2015) and a two-rater system that ensured the raters were blind to subject ID and treatment for that day. Although samples were analyzed for one of four stages of estrous, we clustered phases into either estrus or non-estrus (proestrus, metestrus, diestrus), as described previously (Bakhti-Suroosh et al., 2021; Gaulden et al., 2021; Kerstetter et al., 2008; Peterson et al., 2014; Tan et al., 2019). This clustering allowed for comparison of elevated versus depleted estradiol levels, with estrus showing lower estradiol than the remaining groups (Zachry et al., 2021). Lavages were conducted in the downtime between photometry recordings, but not immediately before testing to minimize the contribution of lavage stress in photometry recordings.

### 2.8. Viral and fiber optic cannula histology

At the conclusion of experiments, rats were anesthetized with a fatal dose of sodium pentobarbital (390 mg/kg, i.p.; Fatal Plus, Covetrus, Portland, ME) and were transcardially perfused with ice-cold phosphate buffered saline (PBS) solution followed by 4% paraformaldehyde (PFA) solution in PBS. Brains were removed and fixed in PFA overnight before being rinsed with PBS and finally put in a 30% sucrose solution for at least 3 days. Using a freezing cryostat, 40-µm thick brain sections were collected and stored at -20°C for long-term storage. dLight expression was assessed following a GFP amplification immunological assay. To amplify the GFP signal from dLight 1.3b, we first washed slices in PBS at room temperature, then incubated in 0.3% PBST (PBS with 0.3% Triton-X), followed by incubation in a blocking solution (0.1% PBST containing 5% Normal Goat Serum (Millipore Sigma, Burlington, MA) and 1% bovine serum albumin (BSA; Millipore Sigma, Burlington, MA)) for one hour at room temperature. Slices were then incubated overnight at 4° C with a chicken anti-GFP polyclonal antibody (1:2000 in 0.1% PBST containing 2.5% Normal Goat Serum and 1% BSA; Catalog # 600-901-215, Thermo Fisher Scientific, Pittsburgh, PA). The next day, slices were washed with PBS at room temperature and were then incubated overnight with a secondary Goat anti-chicken IgY Alexafluor 488 antibody (1:1000 in 0.1% PBST containing 2.5% Normal Goat Serum and 1% BSA; Catalog # A-11039, Thermo Fisher Scientific, Pittsburgh, PA). The next day, slices were washed with PBS at room temperature and then mounted on microscope slides and coverslipped with Vectashield anti-fade mounting medium with DAPI (Vector Laboratories, Plain City, OH). Viral expression and fiber optic cannula placement was then assessed in relation to their position on a Paxinos & Watson brain atlas.

### 2.9. Exclusion Criteria

Male and female rats were excluded from applicable tests if they had incomplete data for a test day, lost catheter patency, lost photometry signal response, or did not show acceptable targeting/viral expression from histology analysis. Because our experimental design was focused on within-subjects differences, we did not include test results that were incomplete (e.g., cocaine trace with Rimonabant treatment but no matched trace with a vehicle pretreatment). Positive photometry signal was assessed prior to IV catheter implantation.

The assessment was completed by examining the dLight 1.3b signal in response to an injection of raclopride (2 mg/kg, i.p.), a D2 receptor antagonist that should elevate dopamine levels in the nucleus accumbens. If the median signal increased above the resting baseline, animals were allowed to continue with testing. Rats with no median signal shift or noticeable increase in dLight transient frequency or amplitude were excluded (data not shown). In total, approximately 15% of rats given catheters lost patency over the course of experiments.

We tested patency with i.v. infusions of propofol (0.2 ml/kg, i.v.) and rats that did not immediately respond were excluded from tests that used i.v. infusions. After behavior, rats were assessed for viral expression and fiber optic cannula placement (Fig. 1 B-D). Overall, we excluded 2 males and 4 females that did not have targeting and expression localized in the NAc_ms_.

**Figure 1.**
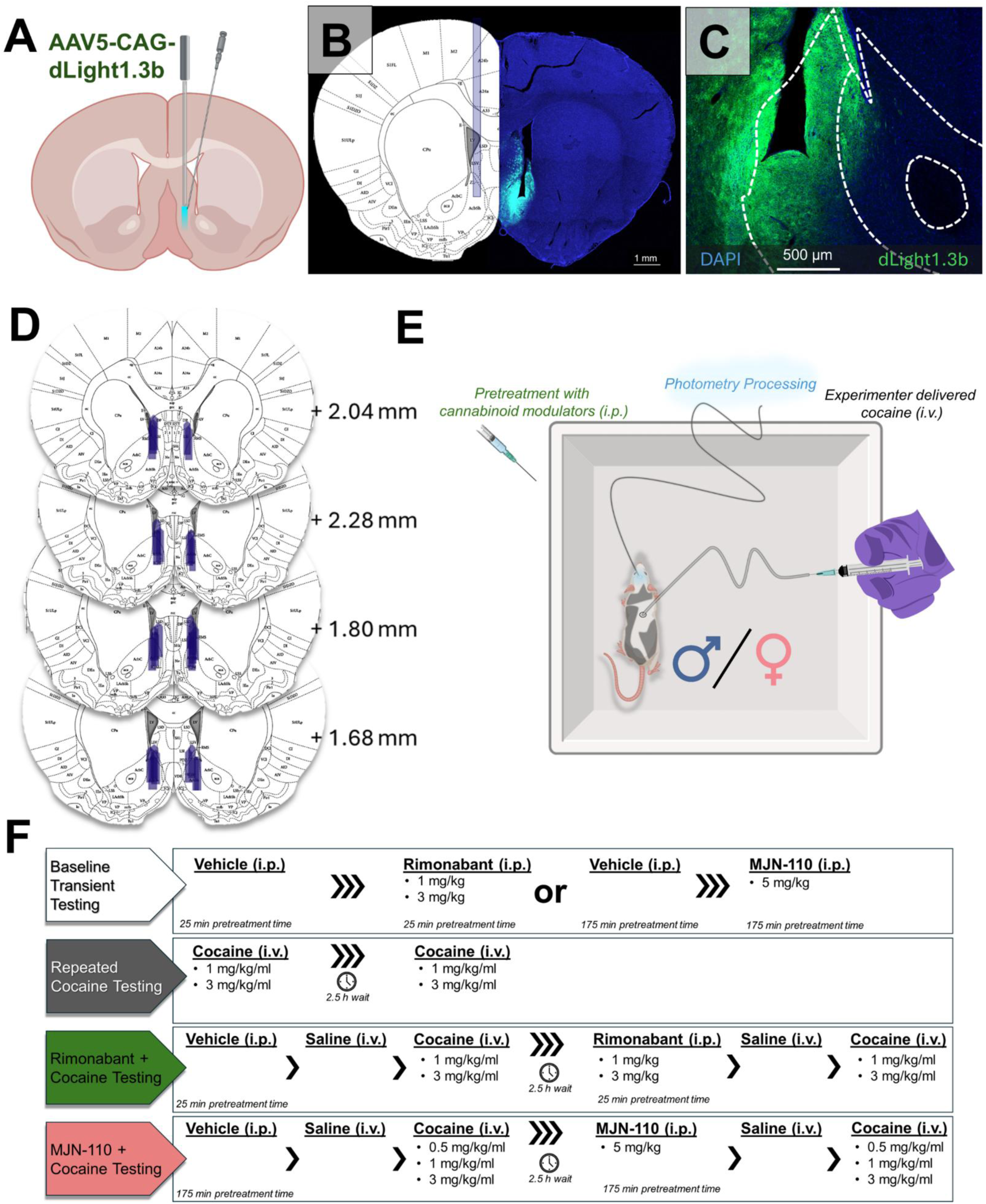
Experimental design for *in vivo* fiber photometry recordings with pharmacological testing. (A) Viral approach and implantation targeting for the NAc_ms_. (B) Representative histology of viral expression and fiber optic cannula targeting. DAPI signal is blue and dLight 1.3b is green. (C) dLight 1.3b expression in the NAc_ms_. Viral expression is prominent around the fiber optic cannula tip, which is situated in the more dorsal aspect of the NAc_ms_. (D) Fiber optic cannula tip placement for all rats included in the study (anterior/posterior distance from *bregma*) (E) Visual representation of behavioral design for fiber photometry experiments. Rats can be pretreated with endocannabinoid-modulating drugs or their corresponding vehicle and tested for photometry signal changes after an experimenter-delivered intravenous infusion of cocaine or saline. (F) Overview of experimental testing regimen. Rats generally completed two or more experimental schedules. For pharmacological testing, vehicle pretreatment always preceded Rimonabant or MJN-110 pretreatment, and the cocaine dose remained consistent for each test day. For cocaine dose response testing, rats received escalating doses of cocaine, starting with a saline control.

### 2.10. Statistical Analysis

Prior to running statistical tests, rats were first subjected to a series of exclusion criteria (detailed above). All tests were conducted after outlier analysis with Grubb’s test, which was set at Q = 1% for detection. Outlier tests, t-tests, 2-way ANOVAS, 3-way ANOVAs, and post-hoc analyses were performed with GraphPad Prism (GraphPad Software Inc., San Diego, CA) or SPSS. For repeated measures designs, any incomplete data was excluded to avoid conducting mixed effects analysis. Post-hoc testing used Holm-Šídák or Dunnett tests for comparing group differences. Given the established data on sex differences in cocaine reward and eCB signaling, we ran statistical tests with the *a priori* hypothesis that males and females would differ in cocaine response, particularly in the context of pharmacological manipulation of eCB signaling. Nonetheless, male and female data were directly compared when applicable, although we argue that our data were not fully powered to uncover all potential interaction effects (i.e. with three-way ANOVAs), particularly because sex differences are generally resolved as magnitude differences and not an inverse relationship. The relevant statistical results are presented in the results section but the full statistical results are reported in tables in the Supplemental Information (Table S1-S14).

### 2.11. Pharmacological Testing Parameters

For at least one day, rats were habituated to the room, behavioral apparatus, and photometry recording equipment that would be used during testing conditions. All pharmacological testing was performed in a small open field apparatus (38.4 cm x 43.8 cm x 35.6 cm length by width by height) with white plastic walls and floor.

All tests were run during the rats’ light cycle (between 11:00 a.m. and 6:00 p.m). Before all photometry recordings, rats were placed in the apparatus with the LEDs turned on for at least five minutes to quench autofluorescence in the fiber optic cable and the brain. To mitigate day-to-day fluctuations in signal quality, pharmacological testing was conducted in a within-subjects schedule in which vehicle injections would occur between 11:00 a.m. and 2:00 p.m., whereas cannabinoid drug testing (i.e. Rimonabant or MJN-110) would occur between 3:00 p.m. and 6:00 p.m. All pharmacological test days were separated by at least 48 hours or more to allow for full washout of all drugs. To minimize order effects, we counterbalanced which dose of cocaine and/or cannabinoid modulator were presented first to animals. For practical reasons, each rat did not complete every possible test (Fig. 1F), but all rats completed at least 2 complete experimental schedules.

Overall, this had a limiting effect on the cumulative cocaine intake for each rat, with each rat receiving no more than 8 total days of cocaine exposure.

### 2.12. Examining the effect of endocannabinoid manipulation on baseline NAcms dopamine transients

Before rats were exposed to cocaine, we tested the effects of endocannabinoid system-targeting drugs on baseline DA transient activity using fiber photometry. For Rimonabant testing, rats were first injected (i.p.) with the 1:1:18 vehicle, and 25 min later were recorded for baseline activity for 10 minutes. Later that day, rats were injected with Rimonabant (1 mg/kg or 3 mg/kg, i.p.), and 25 min later were recorded for baseline activity for 10 minutes. For MJN-110 testing, rats were first injected (i.p.) with the 1:1:8 vehicle, and 175 min later were recorded for baseline activity for 10 minutes. Later that day, rats were injected with MJN-110 (5 mg/kg, i.p.), and 175 min later were recorded for baseline activity for 10 minutes.

### 2.13. Testing the effect of within-day repeat cocaine administration on NAc_ms_ cocaine-evoked dopamine

To test the effects of same-day repeated cocaine infusions on cocaine-evoked dopamine, we conducted photometry recordings in which rats received the same dose of cocaine infusion separated by at least 2.5h. The infusion was a weight-matched dose of cocaine (1 mg/kg/ml or 3 mg/kg/ml) that was infused again at the same dose later that day. For rats that received both repeat cocaine tests, we counterbalanced whether the high dose (3 mg/kg/ml) or moderate dose (1 mg/kg/ml) cocaine test day was presented first. Photometric activity was recorded for one minute before the infusion epoch, and for 10 minutes following the cocaine infusion.

### 2.14. Testing endocannabinoid regulation of cocaine-evoked NAc_ms_ dopamine

To understand the role of eCB signaling in regulating cocaine-evoked NAc_ms_ DA, we conducted fiber photometry recordings of cocaine infusions with or without pharmacological manipulation using the CB1R inverse agonist Rimonabant or the MAGL inhibitor MJN-110. Both cannabinoid drugs were tested with a four-part test design (Fig. 1F), in which photometric activity was tested with 1) saline infusion after pretreatment with vehicle, 2) cocaine infusion after pretreatment with vehicle, then later in the same day 3) saline infusion after pretreatment with cannabinoid drug (Rimonabant or MJN-110) and finally, 4) cocaine infusion after pretreatment with the cannabinoid drug. This four-part test would occur within a day, and with a range of doses for cocaine and/or cannabinoid drug (Fig. 1F), but the cocaine dose was kept the same and repeated for 2) and 4). For all infusions, photometric activity was recorded for a one-minute baseline prior to infusion, and for 5 min after infusion (saline) or 10 min after infusion (cocaine). For Rimonabant testing, we used two doses of Rimonabant (1 mg/kg, 3 mg/kg, i.p.) with a 25 min pretreatment time for two doses of cocaine (1 mg/kg/ml, 3 mg/kg/ml, i.v.). For MJN-110 testing, we used one dose of MJN-110 (5 mg/kg, i.p.) with a 175 min pretreatment time, to allow for accumulation of 2-AG, for three doses of cocaine (0.5 mg/kg/ml, 1 mg/kg/ml, 3 mg/kg/ml, i.v.).

## 3. Results

### 3.1. Male and female rats show similar NAc_ms_ DA responses and locomotor responses to escalating doses of cocaine infusions

For this experiment, male and female rats were given infusions of cocaine on an ascending dose regimen, beginning with saline (Fig. 2A). A two-way repeated measures (RM) ANOVA for cocaine dose (saline, 0.5 mg/kg/ml, 1 mg/kg/ml, 3 mg/kg/ml; i.v.) and sex (male or female) did not show a significant interaction (Fig. 2B; F (3, 51) = 2.46, p > .05, η^2^p=0.13) on AUC for the DA response to cocaine infusions for males (n = 10) or females (n = 9). However, there was a main effect of cocaine dose (F (3, 51) = 43.51, p < .0001, η^2^p=0.72), but no main effect of sex (F (1, 17) = 0.49; p > .05, η^2^p=0.03). A Holm-Šídák’s multiple comparisons test was performed to compare AUC between cocaine dose groups, which revealed that saline AUC was significantly different from all cocaine doses (p < .05), and that all cocaine groups were significantly different from each other (p < .05). The locomotor response to cocaine was also analyzed for males and females together. A one-way RM ANOVA for cocaine doses was significant (Fig. 2C; F (3, 18) = 4.73; p < .05, η^2^p=0.44) for distance traveled, and a Dunnett’s multiple comparisons test revealed significant differences in distance traveled for moderate (1 mg/kg/ml) and high (3mg/kg/ml) doses of cocaine compared to saline (p < .05). A one-way RM ANOVA for cocaine doses was significant (Fig. 2D; F (3, 18) = 4.53; p < .05, η^2^p=0.43) for average velocity, and Dunnett’s multiple comparisons testing revealed significant differences in average velocity for moderate (1 mg/kg/ml) and high (3 mg/kg/ml) doses of cocaine compared to saline (p < .05). These data indicate that male and female rats show a similar dose-response relationship for cocaine-evoked dopamine.

**Figure 2.**
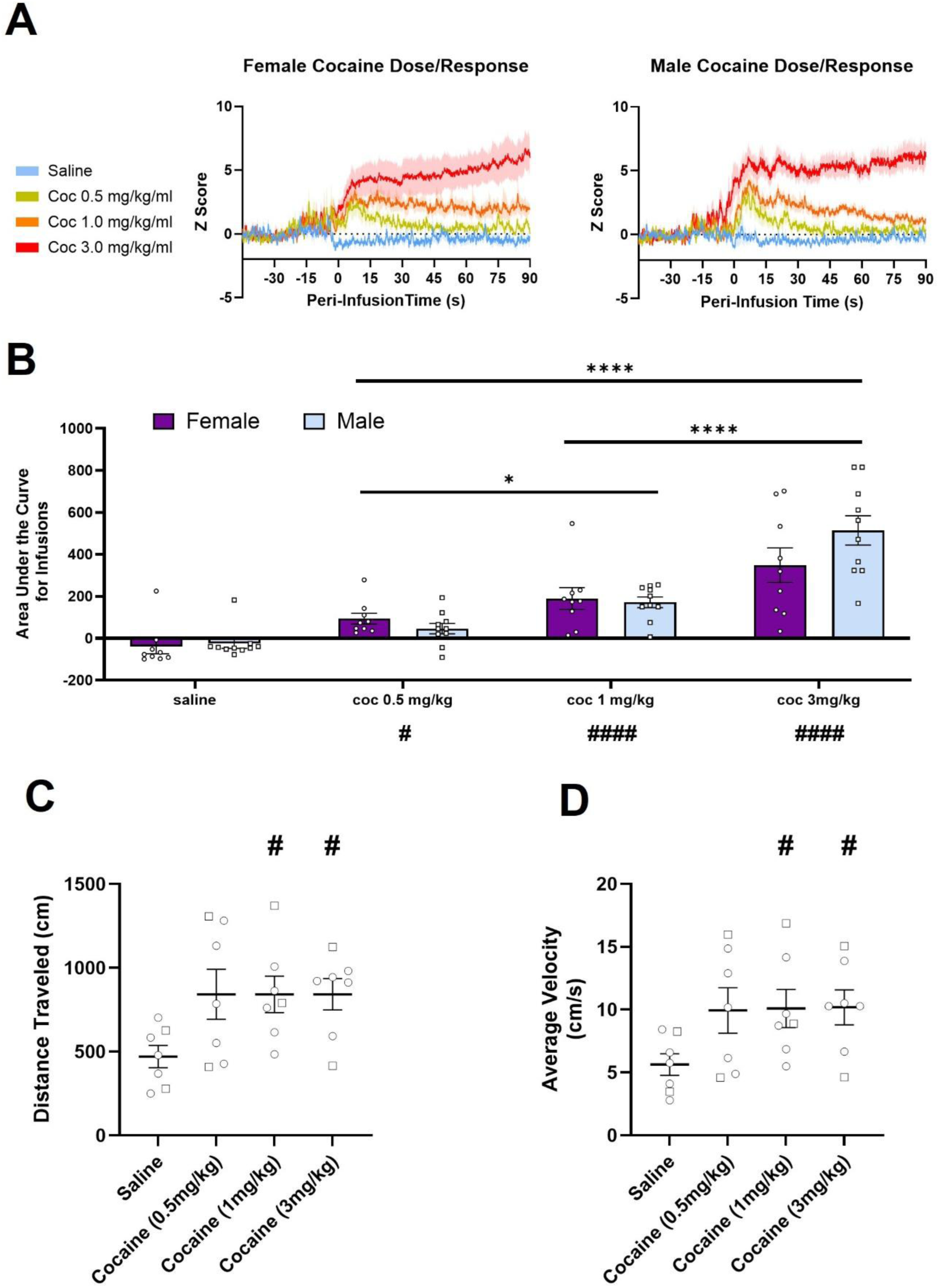
Male and female rats show a consistent dose-response relationship in cocaine-evoked NAc_ms_ dopamine (DA). (A) Z-scored traces of grouped data for males (left, n = 10) and females (right, n = 9) given escalating doses of cocaine. (B) Area under the curve (AUC) analysis for the 90s trace post-infusion. Male and female rats showed similar increases in cocaine-evoked DA from escalating cocaine doses; all cocaine infusions were elevated compared to saline control, and each increase in dose resulted in a significant increase in AUC. (C) Distance travelled in response to infusions over a 5-minute period following iv cocaine administration for male and female rats (n = 7). Moderate (1 mg/kg/ml) and high (3 mg/kg/ml) doses of cocaine significantly increased distance travelled over time compared to saline controls. (D) Average velocity during 5-minute post-infusion period. Rats given a moderate (1 mg/kg/ml) or high (3 mg/kg/ml) dose of cocaine showed significantly elevated average velocity compared to a saline control.

### 3.2. Cannabinoid-targeting drugs do not affect baseline NAc_ms_ DA events

Before exposing rats to cocaine, we first tested the effects of the CB1R inverse agonist Rimonabant and the monoacylglycerol lipase (MAGL) inhibitor MJN-110 on baseline fluorescent dLight 1.3b signal in freely moving male and female rats. We found no effects of pretreatment with either drug on baseline frequency or average event amplitude for dLight 1.3b transient events.

Pretreatment with a low dose of Rimonabant (1 mg/kg, i.p.) did not affect baseline frequency or average amplitude. A two-way repeated measures ANOVA for drug (Vehicle vs Rimonabant 1 mg/kg; i.p.) and sex (male vs female) on dLight event frequency did not reveal a significant interaction (Fig. 3B; F (1, 20) = 1.38, p>.05, η^2^p=0.027) or significant main effects of drug (F (1, 20) = 0.56, p>.05, η^2^p=0.027) or sex (F (1, 20) = 0.25, p>.05, η^2^p=0.012) on dLight event frequency. A two-way repeated measures ANOVA for drug and sex was also conducted for average dLight event amplitude, which did not reveal a significant interaction (Fig. 3C; F (1, 19) = 3.70, p>.05, η^2^p=0.07), nor were there significant main effects for drug (F (1, 19) = 1.21, p>.05, η^2^p=0.06) or sex (F (1, 19) = 0.01, p>.05, η^2^p=0.001) for male (n = 12) and female (n = 11) rats.

**Figure 3.**
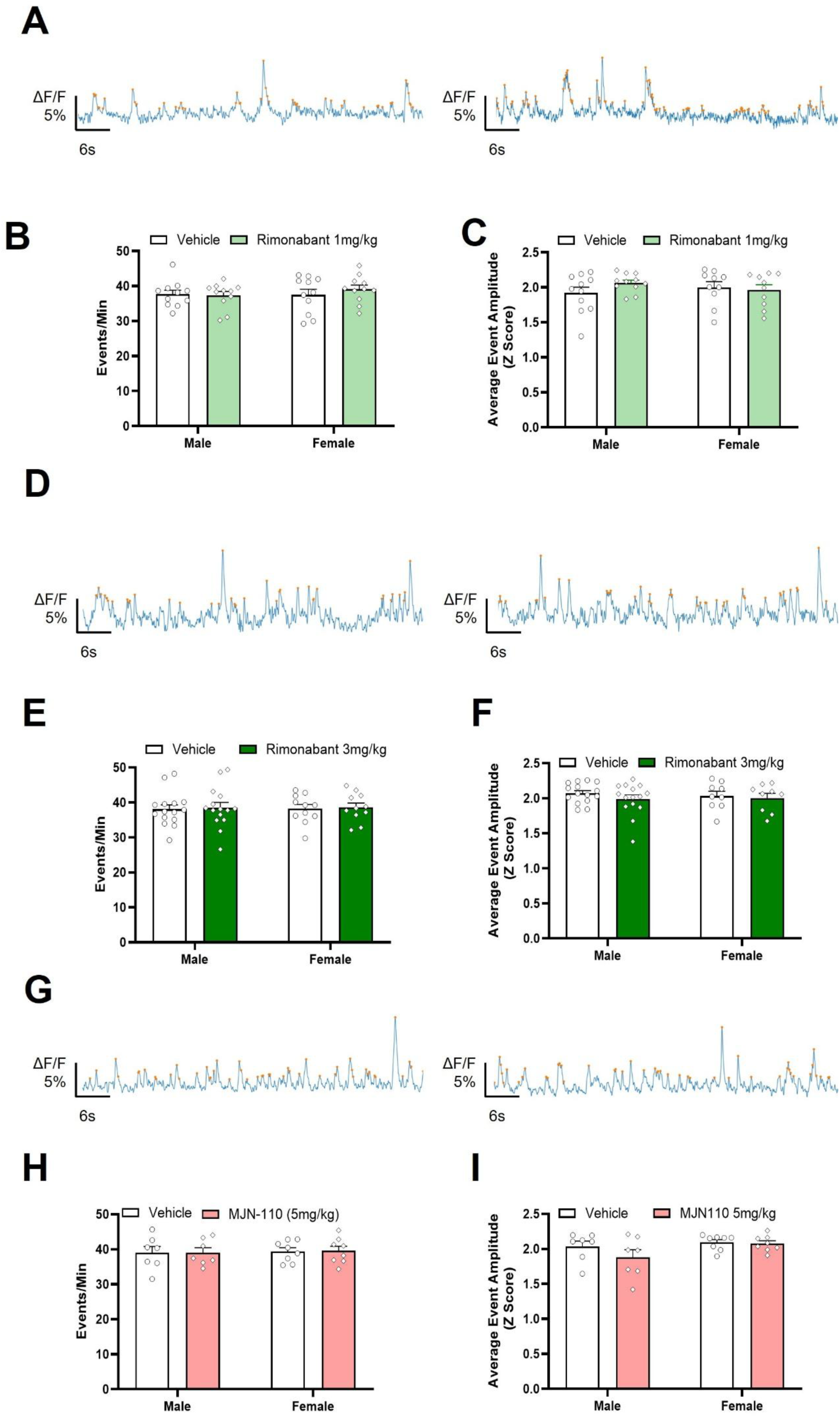
The cannabinoid receptor 1 inverse agonist Rimonabant and MAGL inhibitor MJN-110 did not affect baseline dopamine event frequency or average amplitude. (A) Example traces of baseline transient events for rats treated with either vehicle (i.p.; left) or Rimonabant (1 mg/kg, i.p.; right). (B) A moderate dose of Rimonabant (1 mg/kg, i.p) did not significantly affect baseline event frequency or (C) average amplitude compared to vehicle treatment for males (n = 12) and females (n = 11). (D) Example traces of baseline transient events for rats treated with either vehicle (i.p.; left) or Rimonabant (3 mg/kg, i.p.; right). (E) A higher dose of Rimonabant (3 mg/kg, i.p) did not significantly affect baseline event frequency or (F) average amplitude compared to vehicle treatment for males (n = 16) and females (n= 12). (G) Example traces of baseline transient events for rats treated with either vehicle (i.p.; left) or MJN-110 (5 mg/kg, i.p.; right). (H) MJN-110 (5 mg/kg) treatment did not significantly affect baseline event frequency or (I) average amplitude compared to vehicle treatment for males (n = 8) and females (n = 8).

Pretreatment with a higher dose of Rimonabant (3 mg/kg, i.p.) also did not affect frequency or average amplitude. A two-way repeated measures ANOVA for drug (Vehicle vs Rimonabant 3 mg/kg; i.p.) and sex (male vs female) on dLight event frequency did not reveal a significant interaction (Fig. 3E; F (1, 24) = 0.001, p>.05, η^2^p=0.01) or significant main effects of drug (F (1, 24) = 0.25, p>.05, η^2^p=0.01) or sex (F (1, 24) = 0.01, p>.05, η^2^p=0.01) on dLight event frequency. Similarly, a two-way repeated measures ANOVA did not reveal a significant sex by drug interaction (Fig. 3F; F (1, 22) = 0.37, p>.05, η^2^p=0.017) or significant main effects for drug (F (1, 22) = 1.75, p>.05, η^2^p=0.074), or sex (F (1, 22) = 0.04, p>.05, η^2^p=0.002) on average dLight event amplitude for male (n = 15) or female (n = 11) rats.

Pretreatment with MJN-110 (5 mg/kg, i.p.) also did not affect frequency or average amplitude. A two-way repeated measures ANOVA for drug (Vehicle vs MJN-110 5 mg/kg; i.p.) and sex (male vs female) on dLight event frequency did not reveal a significant interaction (Fig. 3H; F (1, 13) = 0.02, p>.05, η^2^p=0.002) or significant main effects for drug (5 mg/kg, i.p.; F (1, 13) = 0.03, p>.05, η^2^p=0.002) or sex (F (1, 13) = 0.06, p>.05, η^2^p=0.005). A two-way repeated measures ANOVA for average dLight amplitude did not reveal a significant drug by sex interaction (Fig. 3I; F (1, 13) = 2.03, p>.05, η^2^p=0.14) or significant main effects for drug (F (1, 13) = 3.27, p>.05, η^2^p=0.20) or sex (F (1, 13) = 2.27, p>.05, η^2^p=0.15) for males (n = 7) and females (n = 8). Together, these data indicate that treatment with cannabinoid-targeting drugs does not significantly affect baseline NAc_ms_ DA transient features in rats.

### 3.3. Female rats show enhanced sensitivity to CB1R regulation of cocaine-evoked NAc_ms_ DA in response to a moderate dose of cocaine

To understand the contribution of CB1R signaling to the cocaine-evoked NAc_ms_ DA response, we pretreated male and female rats with either vehicle or the CB1R inverse agonist Rimonabant prior to moderate dose (1 mg/kg/ml, i.v.) cocaine infusions. A high dose of Rimonabant was sufficient to attenuate the cocaine-evoked DA response, and this effect was primarily driven by female rats.

A three-way RM ANOVA for drug (vehicle or Rimonabant 1 mg/kg, i.p.), sex (male, n = 8; or female, n = 8), and time (0-30s, 30-60s, 60-90s) did not reveal a significant interaction (Fig 4C,D left; F (2, 28) = 0.68; p > .05, η^2^p=0.05) on AUC, or any two-way interactions or main effects (p > .05). However, a two-way RM ANOVA did reveal a significant drug x sex interaction (Fig 4E, left; F (1, 15) = 4.90; p < .05, η^2^p=0.25) for maximum amplitude across the recording. A Holm-Šídák’s multiple comparisons test did not reveal any significant differences from drug treatment for males or females (p > .05).

**Figure 4.**
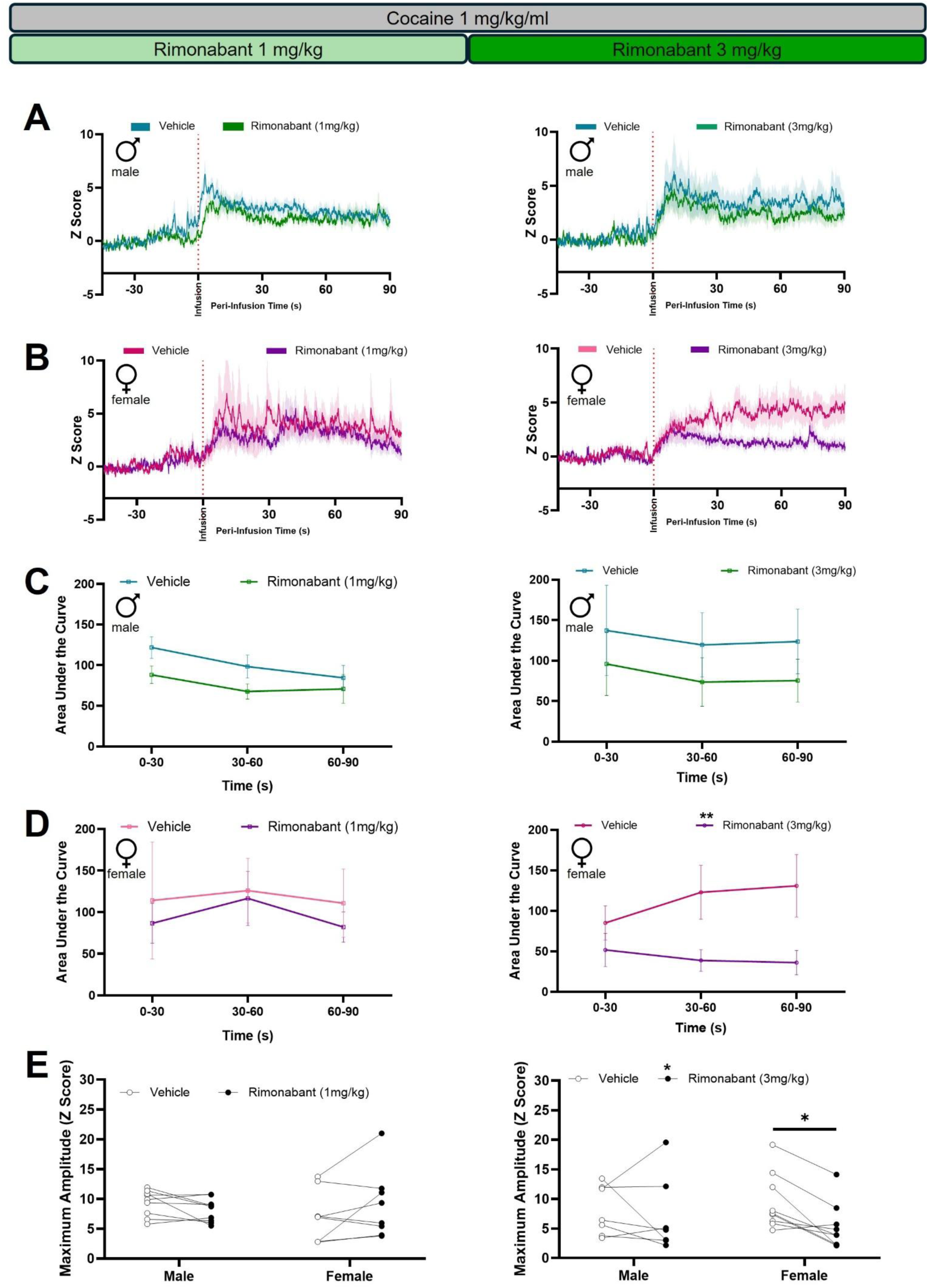
NAc_ms_ DA response to a moderate dose of cocaine (1 mg/kg/ml) is attenuated by a high dose of CB1R inverse agonist Rimonabant, preferentially in female rats. (A) Z-scored average traces for male rats given infusions of a moderate dose (1 mg/kg/ml, i.v.) of cocaine. Male rats were pretreated with vehicle (i.p.) and later in the same day with one of two doses of Rimonabant (1 mg/kg, left; 3 mg/kg, right; i.p.). (B) Z-scored average traces for female rats given infusions of a moderate dose (1 mg/kg/ml, i.v.) of cocaine. Female rats were pretreated with vehicle (i.p.) and later with one of two doses of Rimonabant (1 mg/kg, left; 3 mg/kg, right; i.p.). (C) Area under the curve (AUC) analysis for male NAc_ms_ cocaine-evoked DA response. Pretreatment with either a moderate (n = 8; 1 mg/kg, i.p.; left) or high (n = 8; 3 mg/kg, i.p.; right) dose of Rimonabant did not significantly change cocaine-evoked dopamine as assessed by post-infusion AUC. (D) AUC analysis for female NAc_ms_ cocaine-evoked DA response. Pretreatment with a moderate dose of Rimonabant (n = 8; 1 mg/kg, i.p.; left) did not significantly change cocaine-evoked dopamine as assessed by post-infusion AUC. However, pretreatment with a high dose of Rimonabant (n = 9; 3 mg/kg, i.p.; right) significantly reduced cocaine-evoked dopamine as assessed by post-infusion AUC. (E) Maximum amplitude within the 90s post-infusion trace for male and female rats given either vehicle (i.p.) or Rimonabant. Pretreatment with a moderate dose of Rimonabant (1 mg/kg, i.p.) does not alter maximum amplitude of cocaine-evoked dopamine in male or female rats. (F) Pretreatment with a high dose of Rimonabant (3 mg/kg, i.p.) reduces overall maximum amplitude of cocaine-evoked dopamine, and this effect is primarily driven by female rats. *p<.05, **p<.01, data are presented as mean ± SEM.

A three-way RM ANOVA of drug (vehicle or Rimonabant 3 mg/kg, i.p.), sex (male, n = 8; or female, n = 9), and time (0-30s, 30-60s, 60-90s) revealed a main effect of drug (Fig. 4C,D, right; F (1, 15) = 6.90; p < .05, η^2^p=0.32) but not a significant drug x sex x time interaction (F (2,30) = 0.96; p>.05, η^2^p=0.06) on AUC. We conducted planned two-way ANOVA comparisons based on our *a priori* hypothesis that females show greater sensitivity to Rimonabant treatment than males in the context of cocaine reward (Gaulden et al., 2025). A two-way RM ANOVA did not reveal a significant drug x time interaction for AUC in male rats (F (2, 14) = 0.04; p > .05, η^2^p=0.005), or any significant main effects (p > .05). A two-way RM ANOVA did not reveal a significant drug x time interaction for AUC in female rats (F (2, 16) = 2.02; p > .05, η^2^p=0.20), however, there was a significant main effect of drug (F (1, 8) = 12.22; p < .01, η^2^p=0.60). A two-way RM ANOVA did not reveal a significant sex x drug interaction (Fig. 4E, right; F (1, 14) = 2.26; p > .05, η^2^p=0.14) for maximum amplitude across the recording. However, there was a main effect of drug (F (1, 14) = 5.65; p < .05, η^2^p=0.29). A Holm-Šídák’s multiple comparisons test revealed a significant effect of Rimonabant (3 mg/kg, i.p.) on maximum amplitude in females (p < .05), but not for males (p > .05). In response to a moderate dose of cocaine, a high dose of Rimonabant (3 mg/kg, i.p.) attenuates cocaine-evoked dopamine in female, but not male, rats suggesting that female rats show a greater sensitivity to CB1R regulation of cocaine-evoked dopamine.

### 3.4. CB1R inactivation attenuates cocaine-evoked NAc_ms_ DA in response to a high dose of cocaine, preferentially in female rats

To understand the role of CB1R signaling on cocaine-evoked NAc_ms_ DA in response to a high dose of cocaine, we pretreated male and female rats with either vehicle or the CB1R inverse agonist Rimonabant prior to cocaine (3 mg/kg/ml, i.v.) infusions.

A three-way RM ANOVA of drug (vehicle or Rimonabant 1 mg/kg, i.p.), sex (male, n = 11; or female, n = 12), and time (0-30s, 30-60s, 60-90s) did not reveal a significant three-way interaction (Fig. 5C,D, left; F (2, 42) = 2.99; p > .05, η^2^p=0.13) for AUC, although it was approaching significance (p = .061). However, there was a significant sex by drug interaction F (1, 21) = 4.87; p < .05, η^2^p=0.19) for AUC. In male rats, a two-way RM ANOVA did not reveal a significant time x drug interaction (F (1, 20) = 0.12; p > .05, η^2^p=0.01) for AUC. In females, a two-way RM ANOVA did not reveal a significant time x drug interaction (F (2, 22) = 3.42; p > .05, η^2^p=0.24), although the interaction approached significance (p = .051). However, there were significant main effects for drug (F (1, 11) = 17.59; p < .01, η^2^p=0.62) and time (F (2, 22) = 7.28; p < .01, η^2^p=0.39). A Holm-Šídák’s multiple comparisons test revealed a significant difference from between vehicle and Rimonabant (1 mg/kg, i.p.) across all time points (p < .01). A RM two-way ANOVA revealed a significant sex x drug interaction (Fig. 5E, left; F (1, 22) = 14.34; p < .01, η^2^p=0.39) for maximum amplitude across the post-infusion period. A Holm-Šídák’s multiple comparisons test revealed a significant effect of Rimonabant on maximum amplitude for females (p < .0001) but not for males (p > .05).

**Figure 5.**
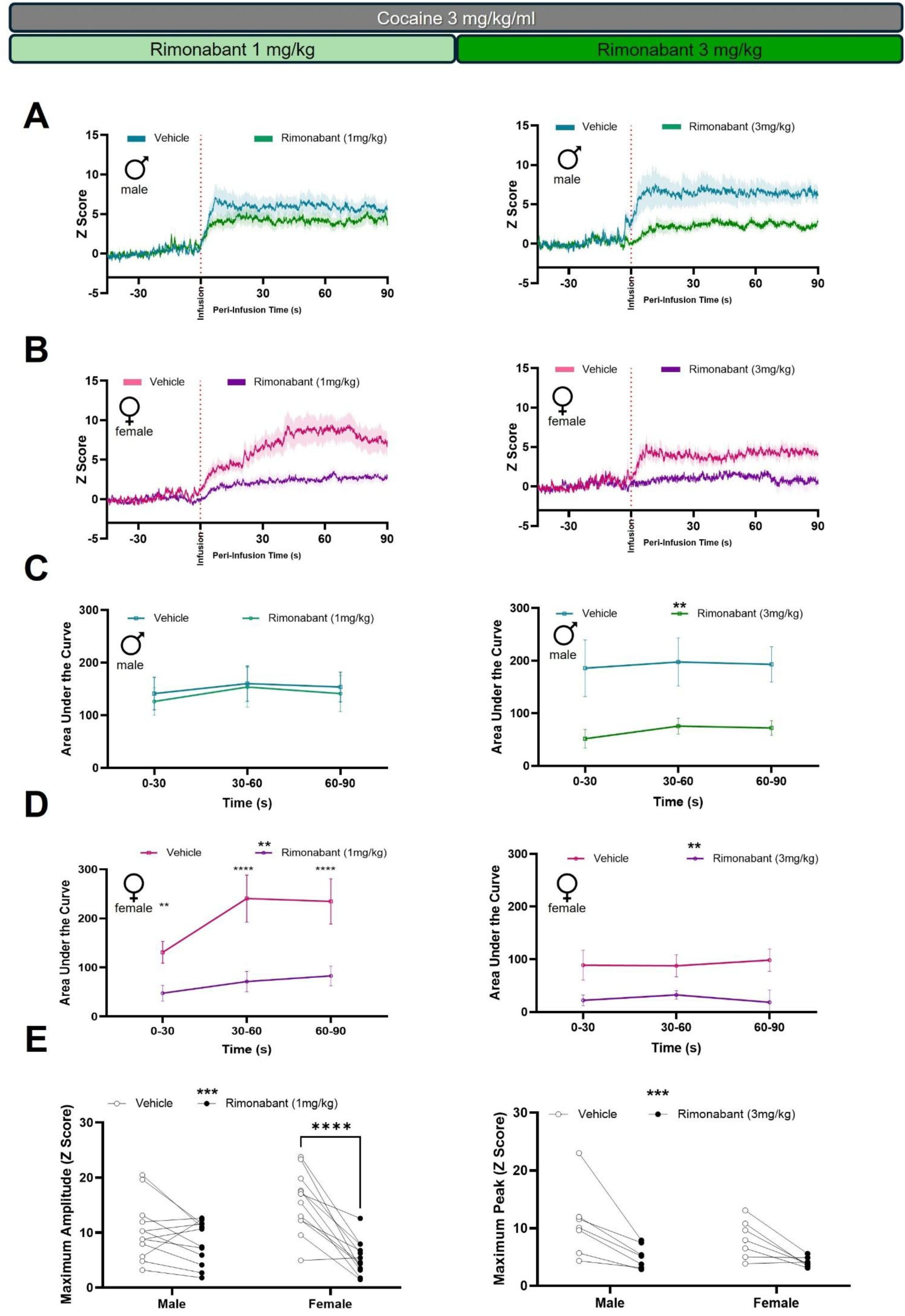
NAc_ms_ DA response to a high dose of cocaine (3 mg/kg/ml) is attenuated by the CB1R inverse agonist Rimonabant, preferentially in females. (A) Z-scored average traces for male rats given infusions of a high dose (3 mg/kg/ml, i.v.) of cocaine. Male rats were pretreated with vehicle (i.p.) and later in the same day with one of two doses of Rimonabant (1 mg/kg, left; 3 mg/kg, right; i.p.). (B) Z-scored average traces for female rats given infusions of a high dose (3 mg/kg/ml, i.v.) of cocaine. Female rats were pretreated with vehicle (i.p.) and later in the same day with one of two doses of Rimonabant (1mg/kg, left; 3 mg/kg, right; i.p.). (C) Area under the curve (AUC) analysis for male NAc_ms_ cocaine-evoked DA response. Pretreatment with a moderate dose of Rimonabant (n = 11; 1 mg/kg, i.p.; left) did not significantly change cocaine-evoked DA as assessed by post-infusion AUC. However, pretreatment with a high dose of Rimonabant (n = 7; 3 mg/kg, i.p.; right) significantly reduced cocaine-evoked DA as assessed by post-infusion AUC. (D) AUC analysis for female NAc_ms_ cocaine-evoked DA response. Pretreatment with either a moderate dose of Rimonabant (n = 12; 1 mg/kg, i.p.; left) or a high dose of Rimonabant (n = 7; 3 mg/kg, i.p.; right) significantly reduced cocaine-evoked DA as assessed by post-infusion AUC. (E) Maximum amplitude within the 90s post-infusion trace for males and females given either vehicle (i.p.) or a moderate dose of Rimonabant (1mg/kg, i.p.). Pretreatment with Rimonabant significantly reduces the maximum amplitude of cocaine-evoked DA in female, but not male rats. (F) Pretreatment with a high dose of Rimonabant (3 mg/kg, i.p.) significantly reduces the maximum amplitude of cocaine-evoked DA in male and female rats. *p<.05, **p<.01, ***p<.001, ****p<.0001, data are presented as mean ± SEM.

A three-way RM ANOVA of drug (vehicle or Rimonabant 3 mg/kg, i.p.), sex (male, n = 7; or female, n = 7), and time (0-30s, 30-60s, 60-90s) did not reveal a significant three-way interaction (Fig. 5C,D right; F (2, 24) = 0.65; p > .05, η^2^p=0.05) or any significant two-way interactions. However, there was a main effect of drug (F (1, 12) = 31.14; p < .0001, η^2^p=0.72) and a trend towards a significant effect of sex (F (1, 12) = 4.67; p = 0.052, η^2^p=0.28). We conducted planned two-way RM ANOVA comparisons based on our *a priori* hypothesis that females show greater sensitivity to Rimonabant treatment than males in the context of cocaine reward (Gaulden et al., 2025). In males, a two-way RM ANOVA did not reveal a significant time x drug interaction (F (2, 12) = 0.29; p > .05, η^2^p=0.05), however there was a significant main effect of drug (F (1, 6) = 17.86; p < .01, η^2^p=0.75). In females, a two-way RM ANOVA did not reveal a significant time x drug interaction (F (2, 12) = 0.92; p > .05, η^2^p=0.13), however there was a significant main effect of drug (F (1, 6) = 14.57; p < .01, η^2^p=0.71). A two-way RM ANOVA did not reveal a significant sex x drug interaction (Fig. 5E, right; F (1, 12) = 0.74; p > .05, η^2^p=0.06) for maximum amplitude in the post-infusion period. However, there was a significant main effect of drug (F (1, 12) = 22.15; p < .001, η^2^p=0.65). A Holm-Šídák’s multiple comparisons test revealed that Rimonabant (3 mg/kg, i.p.) produced a significant decrease in maximum amplitude for both males (p < .01) and females (p < .05). These data indicate that female rats showed greater sensitivity to a moderate dose of Rimonabant, whereas both male and female rats showed similar effects in significant high-dose Rimonabant-induced reductions in cocaine-evoked dopamine in response to a high dose of cocaine.

### 3.5. 2-AG augmentation enhances cocaine-evoked NAc_ms_ DA in response to a low dose of cocaine that is being driven by female rats

To determine the contribution of 2-AG signaling in the context of cocaine-evoked NAc_ms_ DA response, we pretreated male and female rats with either vehicle or the MAGL inhibitor MJN-110 prior to cocaine infusions.

A three-way RM ANOVA of drug (vehicle or MJN-110 5 mg/kg, i.p.), sex (male, n = 8; or female, n = 6), and time (0-30s, 30-60s, 60-90s) revealed a significant three-way interaction (Fig. 6C,D; F (2, 24) = 6.77; p < .01, η^2^p=0.36) for AUC following administration of a low dose of cocaine (0.5 mg/kg/ml, i.v.). There were no other significant interaction or main effects (p > .05). We analyzed male and female rats separately to understand how drug and time effects may differ. In males, a two-way RM ANOVA revealed a significant drug x time interaction (F (2, 14) = 7.83; p < .05, η^2^p=0.53) for AUC. A Holm-Šídák’s multiple comparisons test did not reveal any significant effects of MJN-110 for any time point. In females, a two-way RM ANOVA did not reveal a significant drug x time interaction (F (2, 10) = 2.13; p > .05, η^2^p=0.30) for AUC. However, there was a significant main effect of drug (F (1, 5) = 7.55; p < .05, η^2^p=0.60). A two-way RM ANOVA did not reveal a significant sex x drug interaction (Fig. 6E; F (1, 13) = 0.96; p > .05, η^2^p=0.07) for maximum amplitude across the post-infusion period. These data indicate that elevating levels of 2-AG can enhance cocaine-evoked dopamine in response to a low dose of cocaine and suggests that female rats are more sensitive to this enhancement.

**Figure 6.**
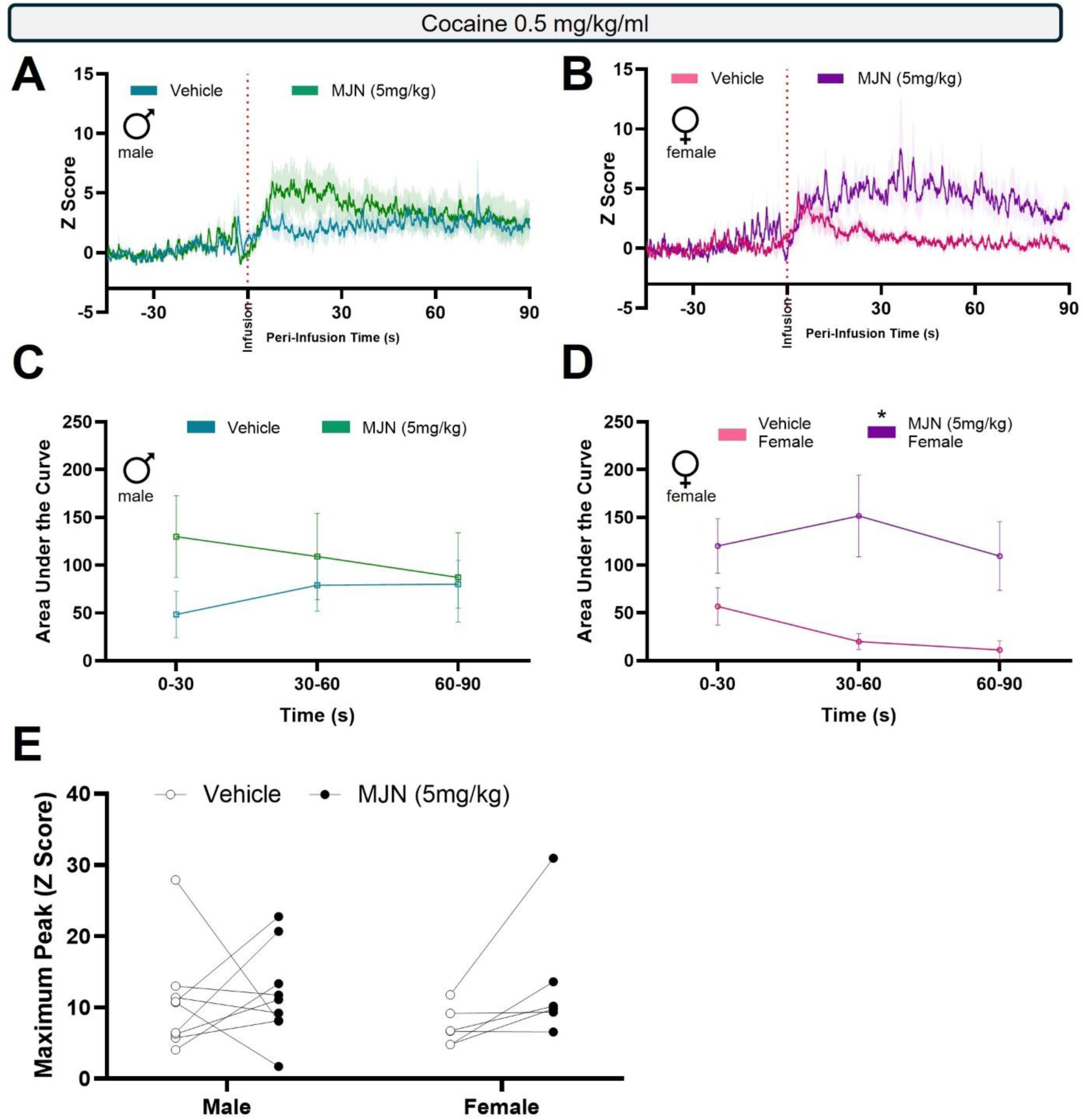
NAc_ms_ DA response to a low dose of cocaine (0.5 mg/kg/ml) is enhanced by the MAGL inhibitor MJN-110, preferentially in female rats. (A) Z-scored average traces for male rats that were given a pretreatment of vehicle of MJN-110 (5 mg/kg, i.p.) followed by an intravenous infusion of a low dose of cocaine (n = 8; 0.5 mg/kg/ml, i.v.). (B) Z-scored average traces for female rats that were given a pretreatment of vehicle of MJN-110 (5 mg/kg, i.p.) followed by an intravenous infusion of a low dose of cocaine (n = 6; 0.5 mg/kg/ml, i.v.). (C) Area under the curve (AUC) analysis for male NAc_ms_ cocaine-evoked DA response. Pretreatment with MJN-110 (5 mg/kg, i.p.) produced transient non-significant increases in cocaine-evoked DA as assessed by post-infusion AUC. (D) AUC analysis for female NAc_ms_ cocaine-evoked DA response. Pretreatment with MJN-110 (5 mg/kg, i.p.) significantly enhanced cocaine-evoked DA as assessed by post-infusion AUC. (E) Maximum amplitude within the 90s post-cocaine infusion trace for male and female rats given either vehicle (i.p.) or MJN-110 (5 mg/kg, i.p.). There were no significant effects of MJN-110 on maximum amplitude of cocaine-evoked DA in male or female rats. *p<.05, data are presented as mean ± SEM.

### 3.6. 2-AG augmentation enhances cocaine-evoked NAc_ms_ DA in response to a moderate dose of cocaine that is being driven by female rats

A three-way RM ANOVA of drug (vehicle or MJN-110 5mg/kg, i.p.), sex (male, n = 7; or female, n = 8), and time (0-30s, 30-60s, 60-90s) did not reveal a significant three-way interaction (Fig. 7C,D; F (2, 26) = 0.30; p > .05, η^2^p=0.02) or any significant two-way interactions for AUC following administration of a moderate dose of cocaine (1 mg/kg/ml, i.v.). However, there was a significant main effect of drug (F (1, 13) = 8.87; p < .05, η^2^p=0.41). Given the observed sex differences in prior experiments, we had an *a priori* hypothesis that females would show greater sensitivity to endocannabinoid manipulation than males. Therefore, we then analyzed the effect of MJN in each sex separately. In males, a RM two-way ANOVA did not reveal a significant drug x time interaction (F (2, 12) = 0.37, p>.05, η^2^p=0.06) nor any significant main effects (p>.05). In females, a two-way RM ANOVA did not reveal a significant drug x time interaction (F (2,14) = 0.80, p>.05, η^2^p=0.10) but did reveal a significant effect of drug (F (1,7) = 6.46, p<.05, η^2^p=0.48). A two-way RM ANOVA did not reveal a significant sex x drug interaction (Fig. 7E; F (1, 13) = 0.19; p > .05, η^2^p=0.02) for maximum amplitude across the post-infusion period. However, there was a significant main effect of drug (F (1, 13) = 5.25; p < .05, η^2^p=0.29).

**Figure 7.**
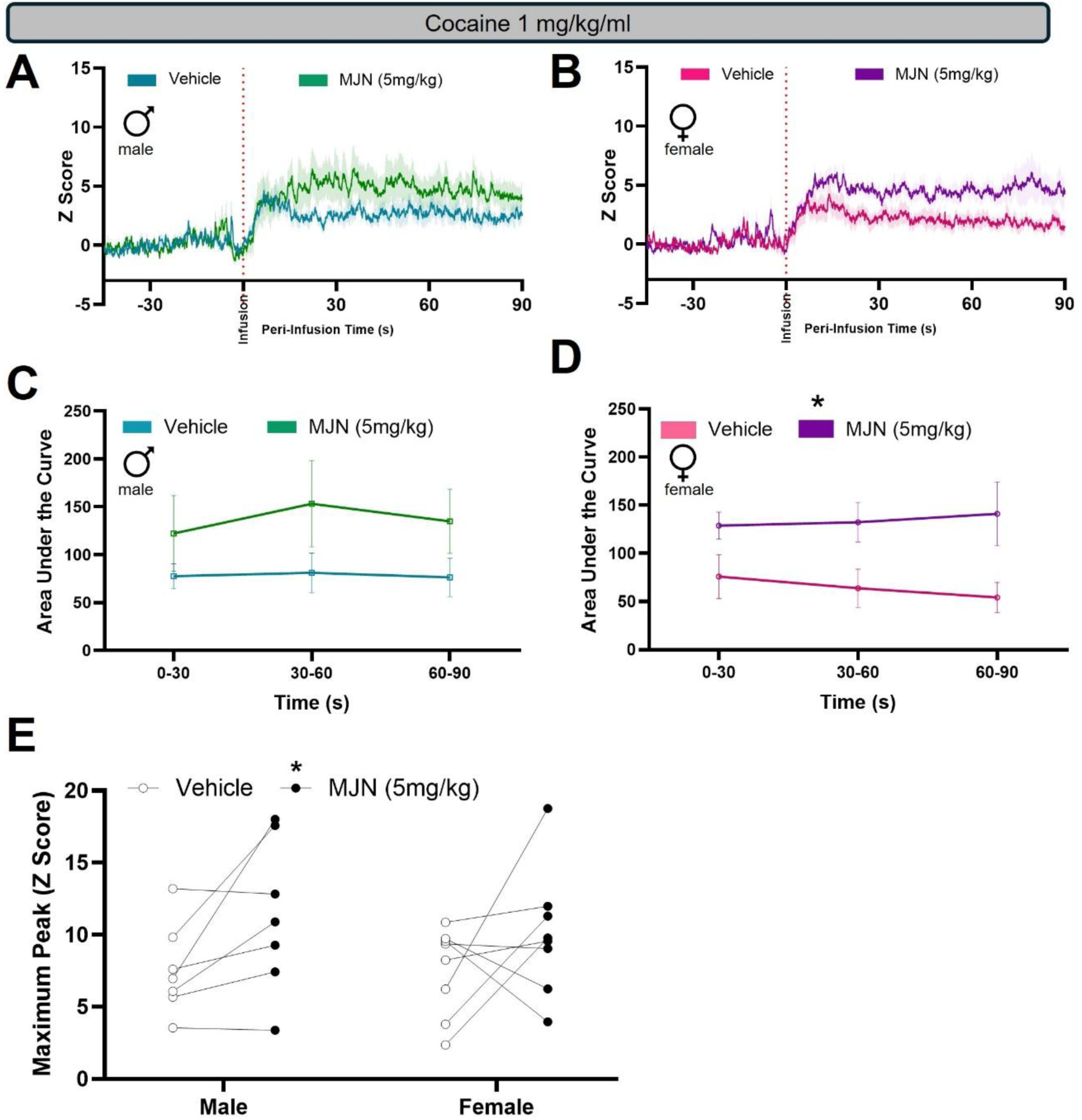
NAc_ms_ DA response to a moderate dose of cocaine (1 mg/kg/ml) is enhanced by the MAGL inhibitor MJN-110, preferentially in female rats. (A) Z-scored average traces for male rats that were given a pretreatment of vehicle of MJN-110 (5 mg/kg, i.p.) followed by an intravenous infusion of a low dose of cocaine (n = 7; 1 mg/kg/ml, i.v.). (B) Z-scored average traces for female rats that were given a pretreatment of vehicle of MJN-110 (5 mg/kg, i.p.) followed by an intravenous infusion of a low dose of cocaine (n = 8; 1 mg/kg/ml, i.v.). (C) Area under the curve (AUC) analysis for male NAc_ms_ cocaine-evoked DA response. Pretreatment with MJN-110 (5mg/kg, i.p.) did not significantly affect cocaine-evoked DA as assessed by post-infusion AUC. (D) AUC analysis for female NAc_ms_ cocaine-evoked DA response. Pretreatment with MJN-110 (5mg/kg, i.p.) significantly enhanced cocaine-evoked DA as assessed by post-infusion AUC. (E) Maximum amplitude within the 90s post-cocaine infusion trace for male and female rats given either vehicle (i.p.) or MJN-110 (5mg/kg, i.p.). MJN-110 pretreatment significantly enhanced the maximum amplitude of cocaine-evoked DA in male and female rats. *p<.05, data are presented as mean ± SEM.

These data indicate that elevation of 2-AG enhances cocaine-evoked dopamine in response to a moderate dose of cocaine and that female rats are primarily driving this effect.

### 3.7. 2-AG augmentation enhances cocaine-evoked female NAc_ms_ DA in response to a high dose of cocaine preferentially in females

A three-way RM ANOVA of drug (vehicle or MJN-110 5 mg/kg, i.p.), sex (male, n = 8; or female, n = 8), and time (0-30s, 30-60s, 60-90s) did not reveal a significant three-way interaction (Fig. 8C,D; F (2, 28) = 0.05; p > .05, η^2^p=0.004) for AUC following a high dose of cocaine (3mg/kg/ml, i.v.). However, there was a significant sex by drug interaction (F (1, 14) = 7.45; p < .05, η^2^p=0.35). In males, a two-way RM ANOVA did not reveal a significant drug x time interaction (F (2, 14) = 0.13; p > .05, η^2^p=0.02) or any main effects for AUC. In females, a two-way RM ANOVA did not reveal a significant drug x time interaction (F (2, 14) = 0.28; p > .05, η^2^p=0.04) for AUC. However, there was a significant main effect of drug (F (1, 7) = 6.22; p < .05, η^2^p=0.47) on AUC. A two-way RM ANOVA revealed a significant sex x drug interaction (Fig. 8E; F (1, 13) = 8.23; p < .05, η^2^p=0.39) for maximum amplitude across the post-infusion period. A Holm-Šídák’s multiple comparisons test did not reveal any significant effects of drug for either sex (p > .05), although females showed a trend towards significant increase from MJN-110 treatment (p = .057). These data indicate that elevation of 2-AG can enhance cocaine-evoked dopamine in response to a high dose of cocaine though the effect is preferentially in female rats.

**Figure 8.**
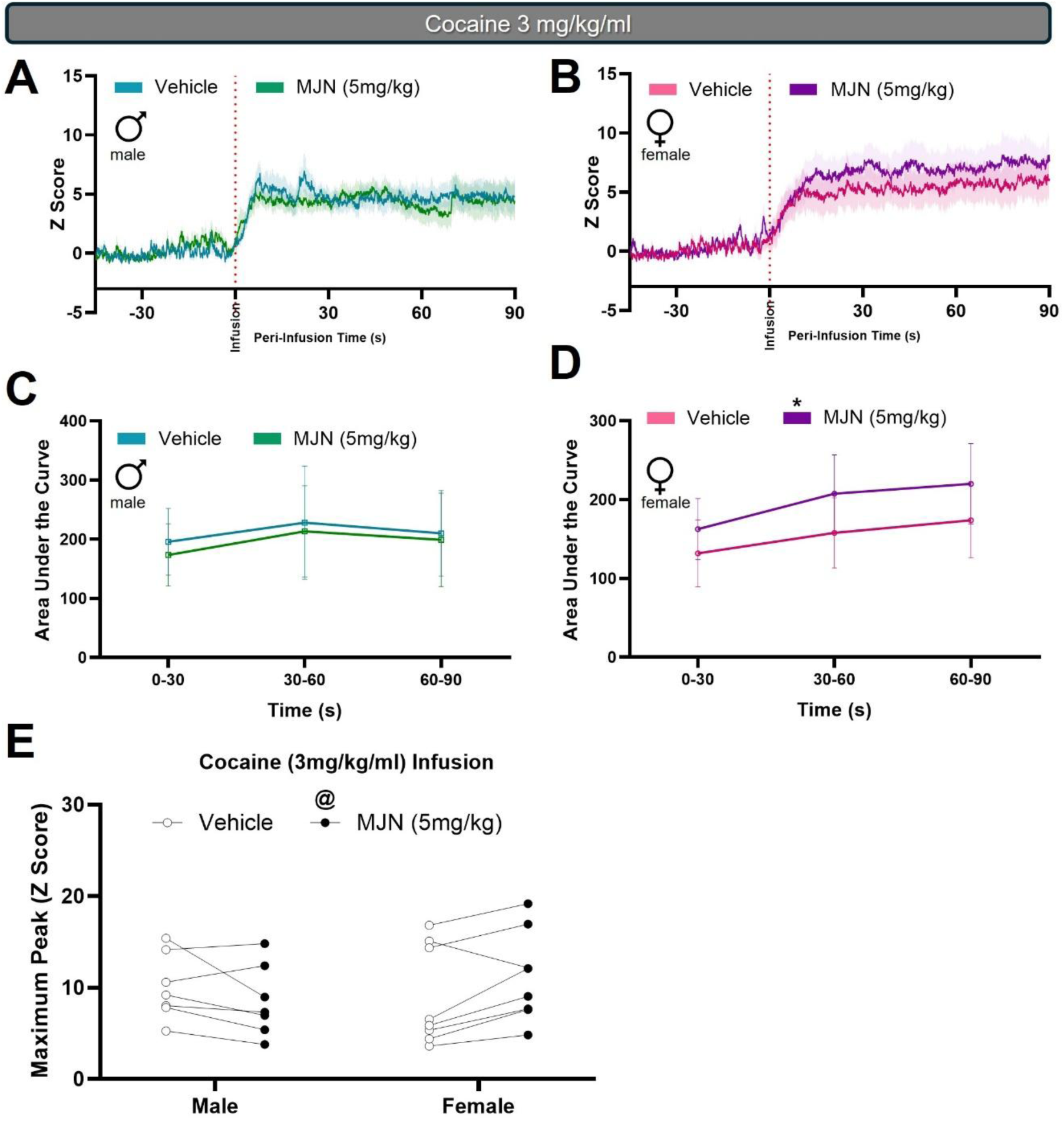
NAc_ms_ DA response to a high dose of cocaine (3 mg/kg/ml) is modestly enhanced by the MAGL inhibitor MJN-110 in female rats. (A) Z-scored average traces for male rats that were given a pretreatment of vehicle of MJN-110 (5 mg/kg, i.p.) followed by an intravenous infusion of a low dose of cocaine (n = 8; 3 mg/kg/ml, i.v.). (B) Z-scored average traces for female rats that were given a pretreatment of vehicle of MJN-110 (5 mg/kg, i.p.) followed by an intravenous infusion of a low dose of cocaine (n = 8; 3 mg/kg/ml, i.v.). (C) Area under the curve (AUC) analysis for male NAc_ms_ cocaine-evoked DA response. Pretreatment with MJN-110 (5 mg/kg, i.p.) did not affect cocaine-evoked DA as assessed by post-infusion AUC. (D) AUC analysis for female NAc_ms_ cocaine-evoked DA response. Pretreatment with MJN-110 (5 mg/kg, i.p.) significantly enhanced cocaine-evoked DA as assessed by post-infusion AUC. (E) Maximum amplitude within the 90s post-cocaine infusion trace for male and female rats given either vehicle (i.p.) or MJN-110 (5 mg/kg, i.p.). There was a significant sex x drug interaction and the females showed a trend towards MJN-110 producing an increase in maximum amplitude of cocaine-evoked DA. *p<.05, @p=.057, data are presented as mean ± SEM.

Locomotor responding during the 90s post-infusion period was analyzed for all experiments involving the CB1R inverse agonist Rimonabant and the MAGL inhibitor MJN-110 (Fig. S1). Locomotor activity is enhanced by cocaine administration in both male and female rats across doses (Table S11). Endocannabinoid modulation modestly reduced overall locomotor activity though it did not impact cocaine-evoked locomotor activity (Table S11).

### 3.8. Repeated cocaine treatment within days, but not across days, produces modest order effects

Our longitudinal design gives rats repeated cocaine infusions across a series of days; therefore, we examined whether the cocaine-evoked dopamine response was altered from the first test in the experiment to the last test for male and female rats (Fig. S2). There were no significant time effects for male or female rats at any dose, indicating that there was no significant sensitization or habituation across the longitudinal study (Table S13).

Because our pharmacological manipulation used cannabinoid-targeting drugs with long half-lives, we always administered vehicle in the morning and cannabinoid-targeting drugs in the afternoon. To identify any potential order effects within a test day, we examined the effects of repeated moderate and high dose cocaine infusions within the same temporal spacing as experiments involving cannabinoid-targeting drugs (Fig. S3). For the moderate dose of cocaine (1 mg/kg/ml), a three-way RM ANOVA for cocaine order (first infusion or second infusion), sex (male, n = 13; or female, n = 12), and time (0-30s, 30-60s, 60-90s) revealed a significant three-way interaction (F (2, 44) = 4.15; p < .05, η^2^p=0.16) for AUC. In male rats, a two-way RM ANOVA revealed a significant cocaine order x time interaction (F (2, 22) = 24.48; p < .001, η^2^p=0.69). A Holm-Šídák’s multiple comparisons test revealed that there was a greater DA response in the 0-30s time bin and a lower DA response in the 60-90s time bin in response to the second infusion of cocaine (p < .05). The repeat infusion of the moderate dose of cocaine (1 mg/kg/ml) alters the time course of the cocaine-evoked DA response in males. In female rats, a two-way RM ANOVA revealed a significant cocaine order x time interaction (F (2, 22) = 11.53; p < .001, η^2^p=0.51) for AUC following a repeated moderate dose of cocaine. A Holm-Šídák’s multiple comparisons test revealed that there was a greater DA response in the 0-30s time bin in response to the second infusion of cocaine indicating a modest sensitization of the cocaine-evoked DA response early following the cocaine infusion in female rats. Together, these data indicate that repeated moderate dose cocaine infusions given over 2h apart produce significantly different patterns of signal response. This is characterized by an initial enhancement of DA signal for the second infusion compared to the first. We also measured the maximum Z score for each infusion. A two-way RM ANOVA for cocaine order and sex did not reveal a significant interaction (F (1, 22) = 0.37; p > .05, η^2^p=0.02) but there was a significant main effect of cocaine order (F (1,22) = 4.72, p < .05), indicating the maximal DA signal is enhanced upon the second infusion of a moderate dose of cocaine within a day.

For repeated administration of a high dose of cocaine (3 mg/kg/ml) within a test day, a three-way RM ANOVA for cocaine order (first infusion or second infusion), sex (male, n = 12; female, n = 10), and time (0-30s, 30-60s, 60-90s) revealed a significant three-way interaction (F (2, 40) = 4.66; p < .05, η^2^p=0.19) for AUC. In males, a two-way RM ANOVA revealed a significant cocaine order x time interaction (F (2, 22) = 11.68; p < .01, η^2^p=0.52) for AUC following administration of a repeated high dose of cocaine. A Holm-Šídák’s multiple comparisons test revealed that there was a lower DA response in the 30-60s and 60-90s time bins in response to the second cocaine infusion of the day compared to the first (p < .05). In females, a two-way RM ANOVA did not reveal a significant cocaine order x time interaction (F (2, 18) = 2.26; p > 05, η^2^p=0.20) nor a main effect of repeated cocaine (F (1, 9) = 2.80, p > .05) for AUC following a repeated high dose of cocaine. Together, these data indicate that repeated high dose cocaine infusions given over 2h apart produce modest attenuation of cocaine-evoked dopamine in response to the second cocaine infusion within a day in male rats but not female rats. We also measured the maximum Z score for each infusion. A two-way RM ANOVA did not reveal a significant cocaine order x sex interaction (F (1, 20) = 1.17; p > .05, η^2^p=0.06), however there was a main effect of cocaine order (F (1, 20) = 5.09, p < .05, η^2^p=0.20), indicating that the maximal amount of DA signal is reduced upon second infusion of a high dose of cocaine.

The cannabinoid-targeting drugs were always associated with the second cocaine infusion of the day and because we have identified some modest effects of repeated cocaine administration within a test day, we assessed whether Rimonabant- or MJN-110-induced changes in cocaine-evoked dopamine were greater than the change observed from repeated cocaine administration alone. We calculated the magnitude of change in the cocaine-evoked DA signal from pretreatment with cannabinoid-targeting drugs vs Vehicle and compared that to the magnitude of change in cocaine-evoked dopamine from repeated cocaine infusions alone (Table S14) within each sex. In males, a two-way RM ANOVA revealed a significant time x treatment interaction (F (2, 40) = 33.96; p < .0001, η^2^p=0.63) for AUC following administration of a moderate dose of cocaine (1 mg/kg/ml) indicating that the Rimonabant (1 mg/kg, i.p.)-induced change in cocaine-evoked DA was greater than that of repeated cocaine administration alone. A similar effect was observed in the females demonstrating a significant time x treatment interaction (F (2, 38) = 8.18; p < .001, η^2^p=0.30). Similarly, a two-way RM ANOVA revealed a significant time x treatment interaction for magnitude change in signal for pretreatment with a higher (3mg/kg) dose of Rimonabant in males (F (2, 40) = 13.58; p < .001, η^2^p=0.40) and females (F (2, 44) = 5.25; p < .05, η^2^p=0.19) compared to repeated moderate dose cocaine infusions alone. Additionally, a two-way RM ANOVA revealed a significant time x treatment interaction for magnitude change in signal for pretreatment with MJN-110 in males (F (2, 38) = 18.45; p < .0001, η^2^p=0.49) and females (F (2, 38) = 12.98; p < .001, η^2^p=0.41) compared to repeated moderate dose cocaine infusions alone. Together, these data indicate that the order effects observed with repeated moderate dose cocaine (1 mg/kg/ml) infusions do not obfuscate the overall magnitude change in cocaine-evoked DA signal from cannabinoid modulation for male and female rats.

We also conducted this comparison for repeated high dose cocaine (3 mg/kg/ml) infusions to compare the magnitude change from cannabinoid modulation versus repeated cocaine alone. A two-way RM ANOVA revealed that there was no significant time x treatment interaction for magnitude change from pretreatment with a moderate dose of Rimonabant (1 mg/kg) prior to administration of high dose cocaine in male rats (F (2, 42) = 0.20; p > .05, η^2^p=0.01) compared to repeated cocaine infusions alone. In females, there was not a significant time x treatment interaction (F (2, 40) = 0.21, p > .05) but there was a significant main effect of Rimonabant (1 mg/kg) treatment (F (2, 40) = 8.18; p < .05, η^2^p=0.24). In males, a two-way RM ANOVA did not reveal a significant time x treatment interaction (F (2, 34) = 3.17; p > .05, η^2^p=0.16) for magnitude change for pre-treatment with a high dose of Rimonabant (3 mg/kg) compared to repeated cocaine infusions alone though it was approaching significance (p = 0.055). However, in females, while there was no significant time x treatment interaction (F (2, 30) = 2.13, p > .05) there was a significant main effect of treatment (F (2, 30) = 2.13; p < .05, η^2^p=0.35) indicating that the Rimonabant (3 mg/kg)-induced attenuation of cocaine-evoked DA was greater than repeated cocaine infusions alone. A two-way RM ANOVA did not reveal a significant time x treatment interaction for magnitude change for pretreatment with MJN-110 in males (F (2, 40) = 2.05; p > .05, η^2^p=0.09) or females (F (1.146, 18.34) = 0.4943; p > .05, η^2^p=0.03) compared to repeated cocaine (3 mg/kg) infusions alone. However, female rats did show a trend towards a significant main effect of drug (p = .057, η^2^p=0.21). These data indicate that the order effects observed with repeated high dose cocaine infusions may obfuscate the overall magnitude change of DA signal from cannabinoid modulation in male rats but not female rats.

### 3.9. Females in estrus show reduced NAc_ms_ cocaine-evoked DA in response to a high dose of cocaine (3 mg/kg/ml)

To examine whether estrous cycle impacts the cocaine-evoked DA response in the *NAc_ms_*, we collected samples of vaginal cytology for stage analysis close to photometry testing of cocaine infusions. Samples were assessed for stage (Fig. 9A) and were then grouped for estrus versus non-estrus classification. We conducted dose-response analysis of cocaine-evoked DA changes using AUC and compared estrus versus non-estrus. A two-way ANOVA for AUC revealed a significant estrous phase x cocaine dose interaction (Fig. 9B; F (2, 53) = 4.20; p < .05, η^2^p=0.14) and a significant main effect of cocaine dose (F (2, 53) = 11.57; p < .0001, η^2^p=0.30), but no significant main effect for estrous phase (F (1, 53) = 0.82; p > .05, η^2^p=0.02). A Holm-Šídák multiple comparisons post-hoc test revealed that in response to the high dose of cocaine (3 mg/kg/ml, i.v.), there was a significant decrease (p < .05) in post-infusion area under the curve for rats in estrus (n = 7) compared to rats in non-estrus phases (n = 16). These data indicate that female rats in estrus display less cocaine-evoked DA in response to a high dose of cocaine compared to those in non-estrus.

**Figure 9.**
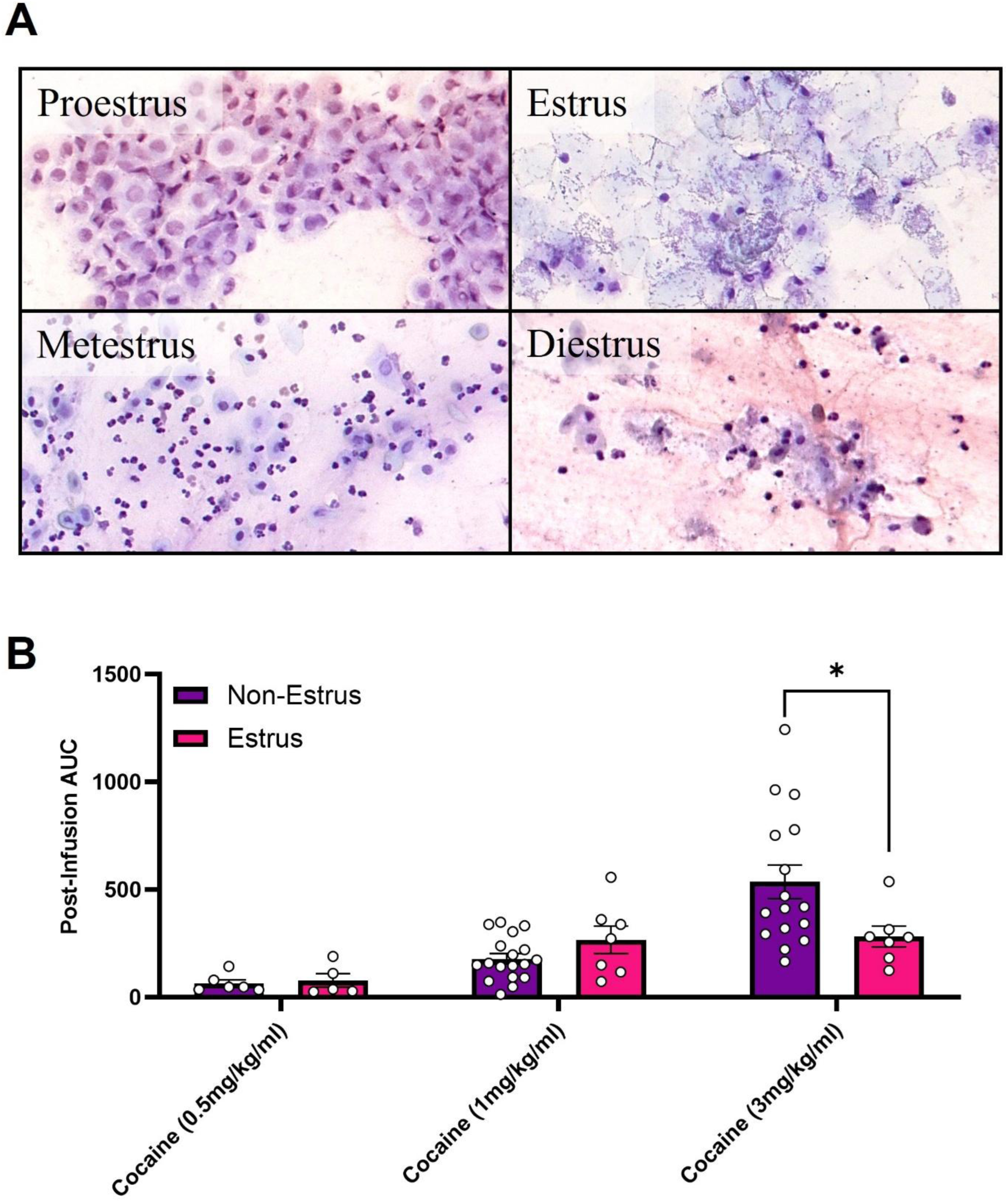
Estrous phase is associated with lower NAc_ms_ cocaine-evoked DA at a high dose of cocaine (3 mg/kg/ml). (A) Representative cytology for estrous cycle staging collected from vaginal lavage samples. (B) Area under the curve analysis of females given different cocaine doses. Stages of the estrous cycle were grouped to compare estrus versus the remaining non-estrus stages. At the high dose of cocaine (3 mg/kg/ml), females in estrus (n = 7) showed significantly reduced cocaine-evoked DA compared to females not in estrus (n = 16; *p < .05 for non-estrus compared to estrus). Data are presented as mean ± SEM.

## 3. Discussion

Others have previously shown that cannabinoid receptor 1 (CB1R) signaling is an important regulator of cocaine-evoked dopamine (DA) release in the NAc_ms_ (Cheer et al., 2007), and that this CB1R regulation is likely localized to the ventral tegmental area (VTA) under baseline conditions (McReynolds et al., 2023; Wang et al., 2015) in male rats. It is critical to understand the factors that can influence this circuit, given the established role of NAc_ms_ DA signaling in regulating cocaine self-administration (SA) behavior (Ito et al., 2004; Pontieri et al., 1995). The data presented here builds upon prior work by using a new technique that enables longitudinal recordings to be able to interrogate the dose-response relationship between CB1R regulation of cocaine-evoked DA. The data also demonstrate that endocannabinoid (eCB) regulation of cocaine-evoked DA is bidirectional with CB1R inactivation attenuating cocaine-evoked DA and 2-AG enhancement elevating cocaine-evoked DA. Importantly, these data examine these relationships in the context of sex as a biological variable and examine endocannabinoid regulation of cocaine-evoked DA in male and female rats. This study reinforces the role of CB1R signaling in regulating cocaine-evoked DA release and supports 2-AG as the principle eCB driving this mechanism. Further, we show sex differences throughout our testing which indicate that females show enhanced sensitivity to eCB modulation of cocaine-evoked DA compared to male rats.

Overall, our results replicate the experiments conducted by Cheer and colleagues (Cheer et al., 2007) who used a fast scan cyclic voltammetry (FSCV) approach. Using similar experimental parameters, but with a fiber photometry approach, we were able to replicate the directionality of CB1R regulation by showing that Rimonabant significantly attenuated cocaine-evoked DA in the NAc_ms_ for males and females in response to a high dose of cocaine (3 mg/kg/ml, i.v.). Although we modeled our experimental design on these previous studies, there were notable differences in our approaches. First, we opted to use higher doses of Rimonabant (1, 3 mg/kg) than what was used in FSCV approaches (0.3 mg/kg). This dose was selected based on prior data from our group and others (Engi et al., 2021; Gaulden et al., 2025; Mereu et al., 2015), which shows robust behavioral responses on food- and drug-motivated behavior. Because fiber photometry is a less sensitive technique compared to FSCV, and because we were interested in behavioral responses to CB1R inverse agonism, we did not include the lower 0.3 mg/kg dose. Our data indicate that male rats would likely not show a significant pharmacological response to a lower Rimonabant dose (0.3 mg/kg), based on the non-significant effects at 1 mg/kg. Second, our approach conducted repeated cocaine testing on a more protracted timescale. To allow more time for cocaine clearance, we delivered the second infusion of cocaine 2.5 h after the first, whereas experiments conducted by Cheer and colleagues (Cheer et al., 2004) delivered cocaine infusions 30 min apart. Third, we leveraged the ability to test subjects multiple times across several weeks with fiber photometry to expand the range of doses for both Rimonabant and cocaine to better understand the relationship between endocannabinoid signaling and cocaine-evoked DA. This also allowed us to measure the cumulative effects of cocaine exposure on the NAc_ms_ circuit, which did not show evidence of sensitization or habituation when several days were given between cocaine administration (Fig. S2). Finally, we also measured locomotor activation following cocaine administration though we did not observe effects of any cannabinoid system modulator on cocaine-induced increases in locomotor activity (Fig S1). This suggests that eCB regulation of NAc_ms_ cocaine-evoked DA may be more important for discrete reward processing which was not examined in the current experimental design. Using these approaches, we showed that a high dose of Rimonabant (3 mg/kg) was sufficient to attenuate the DA response to a moderate dose of cocaine (1 mg/kg/ml) in females and both Rimonabant doses attenuated the DA response to a high dose of cocaine. Females showed a greater sensitivity to CB1R inverse agonism with Rimonabant across tests, suggesting a dose response shift towards greater attenuation of cocaine-evoked dopamine with a lower dose of Rimonabant (1 mg/kg), or more dramatic reduction from Rimonabant at a lower dose of cocaine (1 mg/kg/ml).

Prior research implicates the endocannabinoid 2-AG, but not AEA, in the acute regulation of cocaine-evoked DA release in the NAc_ms_ via *in vitro* studies (Wang et al., 2015). More recently, cocaine treatment has been shown to increase motivation for operant food responding via enhancement of 2-AG signaling in the VTA (Engi et al., 2021). The work described in this study supports these findings by investigating the role of 2-AG signaling on cocaine-evoked NAc_ms_ DA in awake and freely moving rats. While our data implicates 2-AG as the primary eCB mediating these effects, our approach cannot rule out AEA signaling meaningfully contributing to cocaine-evoked DA; AEA has shown to mediate sensitized DA response in the NAc core, but not the NAc shell of mice (Mereu et al., 2015). However, we do show that augmentation of 2-AG via MJN-110, a MAGL inhibitor, is sufficient to enhance cocaine-evoked DA in the NAc_ms_. We selected the 5 mg/kg dose because it has been shown to produce a maximal effect on MAGL inhibition (Niphakis et al., 2013), and we included a lower dose of cocaine (0.5mg/kg/ml) based on the hypothesis that we may hit a “ceiling effect” by augmenting 2-AG signaling prior to a high dose of cocaine (3 mg/kg/ml). This is supported by our data wherein the effects of MJN-110 on cocaine-evoked DA in response to the high dose of cocaine was modest and smaller than the increases observed at lower doses of cocaine. Critically, we show that 2-AG augmentation with MJN-110 increased DA signal for the low, moderate, and high doses of cocaine, with a female-specific effect at the high dose.

However, we did select a dose of MJN-110 that would produce maximal elevation of 2-AG (5 mg/kg; (Wilkerson et al., 2016)) and cannot fully determine whether there is differential sensitivity to 2-AG augmentation of cocaine-evoked DA between sexes. Importantly, while previous studies have implicated 2-AG in the regulation of cocaine-evoked DA, those studies achieved those results by reducing 2-AG levels via DAGL inhibition (Wang et al., 2015). However, here we have demonstrated *in vivo* that enhancing 2-AG levels via MAGL inhibition has the ability to enhance cocaine-evoked DA. This is the first time, to the best of our knowledge, that 2-AG has been shown to directly augment cocaine-evoked dopamine in the NAc_ms_ *in vivo.* These findings support the idea that endocannabinoid signaling can bidirectionally regulate cocaine-evoked dopamine.

Furthermore, our use of a MAGL inhibitor provides activation of CB1Rs as an indirect agonist, an approach that is thought to represent a more physiological enhancement of CB1R signaling as increased activation of CB1Rs should only occur in locations where 2-AG signaling is already occurring. Future work is needed to fully characterize what role AEA might have and how components of the endocannabinoid system might differ for male versus female subjects to understand the differential sensitivity to 2-AG regulation of cocaine-evoked DA between sexes.

The importance of sex as a biological variable has been consistently underscored in clinical literature investigating differences in cocaine reward and cocaine use (Becker & Chartoff, 2018; Becker & Hu, 2008; Fattore et al., 2014). Women generally have a greater propensity for developing cocaine dependence, along with more adverse outcomes from cocaine use (Becker & Koob, 2016). This discrepancy is upheld in preclinical studies of cocaine self-administration, which show enhanced acquisition (Lynch & Carroll, 1999), escalation (Roth & Carroll, 2004), and motivation to use (Roberts et al., 1989) in female rats. Investigation of sexual dimorphism in cocaine reward has shown enhancement of cocaine-evoked DA release in the nucleus accumbens and dorsal striatum of female rodents. For example, Hersey and colleagues have reported that female mice are overall more sensitive to cocaine effects in the enhancement DA release in the NAc shell (Hersey et al., 2023). The work presented here does not support evidence suggesting that female rats have greater overall cocaine-evoked DA release compared to males. However, our methodological approach is narrower in scope than some; we used repeated experimenter-delivered cocaine infusions on a longitudinal study design that did not produce sensitization or habituation (Table S13). Additionally, for practical reasons, we did not study the protracted changes in DA signaling after cocaine infusions (> 90s). These limitations allow us to draw inferences about the underlying neurophysiological differences in cocaine DA responses between male and female rats but not interpret aspects of volitional cocaine use or neuroplastic changes in mesolimbic circuitry over repeated cocaine exposure. However, we do show sex differences in endocannabinoid regulation of cocaine-evoked DA, as well as data indicating that estradiol regulates cocaine-evoked DA. Importantly, females are typically not included in examination of endocannabinoid regulation of cocaine-evoked DA. We have demonstrated here that female rats show increased sensitivity to endocannabinoid regulation of cocaine-evoked DA compared to males highlighting the importance of including females in examination of endocannabinoid regulation of cocaine-related behavior and neuroplastic changes. Additionally, this increase in sensitivity is in line with prior work from our lab that identified that female rats are more sensitive to CB1R regulation of cocaine self-administration behavior (Gaulden et al., 2025). Taken together, better understanding of how these effects observed in females corresponds to regulation of cocaine-related behaviors needs to be further explored.

Given the obvious contrasts in circulating gonadal hormones (e.g., estradiol, progesterone) between males and females, understanding the role of gonadal hormones in the regulation of cocaine reward has been a main focus for preclinical sex differences research. Although male gonadal hormones fluctuate, and can impact mesolimbic function (Holloway & Lerner, 2024), they have not been shown to modulate cocaine self-administration for males (Jackson et al., 2006). In contrast, females show hormonal regulation of cocaine reward and cocaine-seeking behavior. Progesterone, which peaks and declines just before the luteal phase in rats, and during the luteal phase in humans, has been shown to be protective against cocaine craving in rodents (Doncheck et al., 2020; Zlebnik & Cheer, 2016) and humans (Milivojevic & Sinha, 2018; Moran-Santa Maria et al., 2018) but does not seem to modulate the rewarding effects of cocaine (Peart et al., 2022).

Conversely, estradiol has been associated with enhancing cocaine-induced dopaminergic signaling within the NAc and dorsolateral striatum of female rodents (Cummings et al., 2014) by enhancing cocaine’s affinity for the dopamine transporter (DAT; Calipari et al., 2017) and by upregulating cocaine-evoked dopamine release (Yoest et al., 2018). Estradiol increases VTA DA neuron activity in part by binding to estrogen receptor alpha (ERα) located on inhibitory inputs from the medial preoptic area (mPOA). ERα binding silences mPOA inhibitory inputs to VTA DA neurons (McHenry et al., 2017), allowing more basal DA signaling, as well as enhancing cocaine reward (Robison et al., 2018). Further, estradiol rapidly reduces the binding efficiency of D2 autoreceptors in the striatum, effectively reducing the suppression of VTA DA release (Bazzett & Becker, 1994). Finally, activation of estrogen receptor beta (ERβ) potentiates cocaine-evoked DA release (Satta et al., 2018; Yoest et al., 2018), likely via VTA activation (Creutz & Kritzer, 2002).

The data presented in this study supports the body of evidence suggesting that cocaine-evoked VTA DA activity should be lowest when estradiol is lowest (i.e., in the estrus stage). Our experiments were not powered to detect differences between all stages of the estrous cycle; instead we followed grouping conventions used previously by our group and by others (Bakhti-Suroosh et al., 2021; Gaulden et al., 2021; Kerstetter et al., 2008; Peterson et al., 2014; Tan et al., 2019) to contrast low estradiol levels found in estrus with the moderate to high levels of estradiol found in the remaining stages. Using this methodology, we show that for female rats given the highest dose of cocaine (3 mg/kg/ml), those in estrus have a reduced overall cocaine-evoked DA response. It is likely that we only show significant effects of estrus versus non-estrus at this highest dose because of a “floor” effect at low and moderate cocaine doses which can obfuscate DA reduction in estrus. Additionally, there are inherent challenges of collecting a sufficient sample size for females in any one stage of the estrous cycle. Therefore, we posit estradiol is also a mediator of cocaine-evoked DA at low and moderate doses of cocaine, although our ability to demonstrate this effect is challenged by the sensitivity of the technique used as well as sample size.

Throughout this study, female rats showed increases in sensitivity to CB1R modulation, which could indicate sex differences in eCB mechanisms. Although literature describing sex differences in the eCB system is somewhat sparse, there is evidence that females show more tonic 2-AG signaling that regulates the basal activity of VTA DA neurons (Melis et al., 2013), and that estradiol can indirectly promote the production of eCBs to drive cocaine reward (Peterson et al., 2016). Our experimental design included analysis plans for investigating the role of estrous phase in CB1R modulation of cocaine-evoked NAc_ms_ dopamine; however, our sample size was underpowered to detect these differences (data not shown). Our future plans include accumulating a sufficiently powered dataset to draw inferences about how estrous phase or circulating gonadal hormones might influence cannabinoid regulation of cocaine reward. Although our data was not statistically powered for detection of estrous cycle during 2-AG augmentation of cocaine infusions, we do show evidence that females overall were more sensitive to 2-AG augmentation than males at the moderate and high dose cocaine infusions. Furthermore, females showed more of sustained enhancement of low dose cocaine-evoked DA following MAGL inhibition compared to the more transient effect observed in the males. This may indicate that female rats have upregulated production of 2-AG synthesis in the mesolimbic DA pathway or that there may be sex differences in clearance and degradation of 2-AG that is MAGL-independent. However, understanding whether this effect is mediated by estrous cycle, and how estradiol might regulate 2-AG production (Bálint et al., 2016; Peterson et al., 2016) warrants future investigation.

Fiber photometry has technical advantages over other DA recording techniques, such as FSCV and microdialysis, because of its accessibility, consistent quality over repeated uses, and relatively fast sample rate (Dong et al., 2025). Although fluorescent activation via DA biosensors cannot directly measure sub-second DA dynamics like FSCV, the binding kinetics of dLight 1.3b are relatively fast, particularly on the timescale of cocaine-evoked DA dynamics. Additionally, the dLight family of biosensors is resistant to internalization and maintains lability during consistent agonism (Leopold et al., 2019; Patriarchi et al., 2020). However, cocaine treatment boosted the median signal towards a ceiling effect and likely obfuscated individual DA events, particularly in our Z-scored data. This is a technical challenge for many pharmacology approaches to DA biosensors (Wallace et al., 2025). For our purposes, cocaine-induced bulk signal changes were the primary data of interest. Given the relatively narrow dynamic window for photometry recordings with the current DA biosensors, we argue that the reported effects of endocannabinoid modulation and sex differences are robust. However, we acknowledge that we are unable to provide data for how individual DA event occurrences are modulated by our manipulations. Future studies that value individual DA event metrics during cocaine exposure will require a more careful analysis, particularly if using fiber photometry.

Our overall experimental design leveraged the repeatability of fiber photometry testing to uncover the effects of eCB system manipulation on cocaine-evoked DA. To limit the impact of day to day fluctuations in fiber photometry recording quality and to make reliable determinations about how estrous cycle affects these manipulations, we conducted recordings comparing vehicle versus Rimonabant or MJN-110 pretreatment within the same day. This required repeated cocaine infusions to occur and also required vehicle treatment to always precede cannabinoid modulators, which, based on their half-life could be as short as 2 h (Järbe et al., 2010) or as long as 15 h (McLaughlin et al., 2003) for systemic Rimonabant, and a purported half-life of 3 h for MJN-110 (Ma, 2022). Given these constraints, we acknowledge that our results are challenged by order effects from cannabinoid pharmacological manipulation always corresponding to the second cocaine infusion of the day and always occurring later in the afternoon. To address this limitation, we conducted control experiments in which we delivered repeated cocaine infusions with the same doses and time constraints as our pharmacological testing of cocaine infusions.

We were surprised by the results of within-day repeated cocaine infusions (Fig. S3); both male and female rats showed an initial increase in cocaine-evoked dopamine on the second infusion of the day at the moderate dose of cocaine (1 mg/kg/ml), and a decrease in cocaine-evoked dopamine on the second infusion of the day at the high dose of cocaine (3 mg/kg/ml), although this effect was likely predominately driven by males. This initial increase in cocaine-evoked DA from a repeated moderate dose of cocaine is indicative of sensitization, which occurs within 2h of an initial moderate dose infusion of cocaine (1 mg/kg), as previously demonstrated with FSCV recordings in the NAc shell of male rats (Singer et al., 2017). However, our fiber photometry data indicates that while there is an initial increase in cocaine-evoked DA from repeated infusions of a moderate dose of cocaine, this difference only occurs within the initial ∼30s after the infusion and is compensated by a more pronounced decline in cocaine-evoked DA after the initial peak, at least in males. This change in temporal dynamics is not shown in FSCV recordings (Singer et al., 2017), and more work is needed to determine whether this contrast is attributed to differential sensitivity or kinetics between FSCV and fiber photometry techniques. We also show a robust increase in locomotor activation from repeated infusions of a moderate dose of cocaine, which is more congruent with cocaine-induced sensitization of mesolimbic reward neurocircuitry, as reported by others (Argilli et al., 2008). There is also evidence that a single high dose injection of cocaine produces a cannabinoid-dependent sensitized locomotor response 24h later that is associated with increased DA release in NAc core, but not NAc shell, from subsequent cocaine infusions in male mice (Mereu et al., 2015). In line with these findings, we showed reduced NAc shell DA following repeated infusions of a high dose of cocaine within the same day, indicating a potential tolerance effect that appears to be driven primarily by male rats. If these dynamics are maintained during pharmacological testing with Rimonabant and MJN-110, we expect that Rimonabant effects may be occluded or obscured by expected decreases from repeated infusions of a high dose of cocaine, whereas MJN-110 effects may be occluded or obscured by initial increases in DA following repeated infusions of a moderate dose of cocaine. Although we cannot rule out this interaction, our within day magnitude change analysis (Table S14) for both drugs shows that pharmacological changes are more robust than the effects of repeated cocaine infusions alone. In contrast to a shift in DA dynamics from high and moderate doses of repeated cocaine, we also show that distance traveled increased from repeated infusions of a high dose of cocaine, indicating behavioral sensitization, although perhaps not as strongly as with a moderate dose of cocaine because of a maximal limit on locomotor activity. Because we show overall acute behavioral sensitization from repeated cocaine infusions, and because the NAc_ms_ is a known regulator of behavioral sensitization (Todtenkopf et al., 2002), more work is needed to explain the disparity between our DA recordings, locomotor activity, and existing FSCV data. To this point, some potential explanations are detailed below.

The mechanism for the decrease in cocaine-evoked DA observed following repeated infusions of a high dose of cocaine may be the decreased efficiency of cocaine to bind to and inhibit DAT following repeated cocaine exposure (Ferris et al., 2011; Siciliano et al., 2016). However, this control experiment was conducted at the onset of testing with drug-naïve rats, therefore the cumulative effects of cocaine exposure were minimized. Adding to this complexity, others report delayed kinetics in the DA response to the first exposure to cocaine (Stuber et al., 2003), as well as delayed NAc_ms_ DA release and clearance for females (Kuiper et al., 2024), and we did qualitatively notice these phenomena for some recordings. Regardless, our experimental paradigm was designed to avoid any cumulative effects of cocaine by spacing out cocaine exposure days by 24 hours or more, and only administering two cocaine infusions, except in the case of the final cocaine dose response experiments. Further, we analyzed dose-matched cocaine responses from the first exposure to the last exposure across the weeks of fiber photometry recordings for each rat and found no significant differences for any dose (Fig. S2) indicating that there was no overall sensitization of the cocaine-evoked DA response across days. Together, the results of the control experiments allowed us to unmask the effects of repeated cocaine treatment with or without cannabinoid modulation by calculating magnitude changes. These comparisons show a clear female-specific sensitivity to CB1R inverse agonism with Rimonabant and 2-AG augmentation with MJN-110.

The work presented here demonstrates a clear role for sex and eCB signaling in the regulation of cocaine-evoked DA in the NAc_ms_. However, there are some additional considerations when interpreting these results. First, cocaine was delivered via experimenter-delivered infusions. This non-volitional design does not model the motivational salience of rodent self-administration models, nor does it measure the changes in mesolimbic function (Siciliano et al., 2015; Siciliano et al., 2016; Willuhn et al., 2012) over the course self-administration behavior. Second, although we show that augmenting 2-AG is sufficient to elevate cocaine-evoked DA, we did not conduct experiments that would test the necessity of 2-AG in the cocaine-evoked DA response (e.g., treatment with a DAGL inhibitor). Based on our results blocking the activation of CB1Rs with Rimonabant, and existing data showing the role of 2-AG in cocaine-evoked DA signaling (Wang et al., 2015) *in vitro*, we hypothesize that 2-AG signaling comprises a significant portion of the cocaine-evoked DA response shown here. Third, our magnitude change analysis of repeated cocaine infusions versus repeated cocaine infusions with a cannabinoid modulator suggest that the attenuation of repeated infusions of a high dose of cocaine obfuscates the effects of Rimonabant, particularly in male rats. However, it is important to also consider that the repeated cocaine alone control experiments were not subject matched to the cannabinoid-modulated cocaine infusions. Therefore, there could be a degree of variability from cohort differences that is difficult to account for. Together, more work is needed to determine how the mechanisms described here integrate into our current understanding of cocaine use and other relevant cocaine-related behaviors. However, there is significant evidence supporting the role of eCB signaling and sex in cocaine use in human and rodent studies.

Individuals who use cocaine likely have disrupted endocannabinoid signaling (Elliott et al., 2025; Tanda, 2007). Specifically, individuals with cocaine dependence have increased plasma 2-AG levels compared to recreational users or controls (Kroll et al., 2023). This change is positively correlated with mGluR5 binding in the caudate, indicating a mechanism by which cocaine use can upregulate circulating 2-AG. On the other hand, a two-week abstinence period for individuals with cocaine use disorder (CUD) is associated with decreased 2-AG, but increased AEA content (Pavón et al., 2013). Because cocaine use and CUD are complex psychosocial constructs, it is important to understand how contributing factors such as stress and sex might influence eCB regulation. For example, cocaine users show elevated glucocorticoids in hair, which positively correlates with endocannabinoid content and level of cocaine use (Voegel et al., 2022). Rodent models of cocaine use clearly identify stress as a contributing factor to cocaine use (Mantsch et al., 2007) that is endocannabinoid-dependent (Gaulden et al., 2025; McReynolds et al., 2023). Adding to this, females show enhanced CB1R regulation of reward-related behaviors, as evidenced by female-specific sensitivity to Rimonabant under non-stress conditions (Bacharach et al., 2018; Gaulden et al., 2025). This sex effect could arise from differences in CB1R signaling in the mesolimbic DA system (Bacharach et al., 2023; Melis et al., 2013; Peterson et al., 2016). Females show slightly decreased overall striatal CB1R affinity (de Fonseca et al., 1994), estradiol-induced eCB production (Peterson et al., 2016), and enhanced sensitivity to cannabinoid drugs when estradiol is elevated (Kim et al.; Salemme et al., 2024) in preclinical rodent models. Further, because women who use cocaine are more at risk for escalating use and developing dependence (Becker & Hu, 2008), therapeutic approaches that target eCB signaling may be particularly effective for mitigating cocaine use patterns in women. In light of the data presented here, future work investigating the efficacy of targeted attenuation of 2-AG (e.g., through use of DAGL inhibitors) on cocaine reward may show better efficacy for female subjects.

## 5. Conclusion

The present study provides clear evidence that endocannabinoid signaling bidirectionally modulates cocaine-evoked dopamine in the nucleus accumbens medial shell, and that this regulation is influenced by biological sex. We demonstrate that CB1R inactivation attenuates, and 2-AG augmentation enhances, cocaine-evoked dopamine *in vivo*, with female rats displaying greater sensitivity to both manipulations. This female-specific sensitivity was further supported by estrous cycle-dependent modulation of cocaine-evoked dopamine, suggesting a role for circulating gonadal hormones, particularly estradiol, in shaping endocannabinoid-dopamine interactions. Together, these data underscore sex as a critical variable in the neuropharmacology of cocaine reward and suggest that endocannabinoid signaling, particularly 2-AG-mediated CB1R activation, may contribute to sex differences in vulnerability to cocaine use disorder. These findings support the development of sex-informed, endocannabinoid-targeted interventions as potential therapeutic strategies for cocaine addiction.

## Data Statement

Data will be publicly available upon publication of the manuscript through University of Cincinnati UCFigshare repository.

## Author Contribution

Jayme McReynolds: Conceptualization, Methodology, Data Analysis, Writing – original draft, reviewing, and editing, Supervision, Funding acquisition. Andrew Gaulden: Conceptualization, Methodology, Investigation, Data analysis, Writing – original draft, reviewing, and editing, Funding acquisition. Kristin Chase: Investigation, Data analysis

