## Supplemental Information for "Sex-specific differences in endocannabinoid regulation of cocaine-evoked dopamine in the medial nucleus accumbens shell"

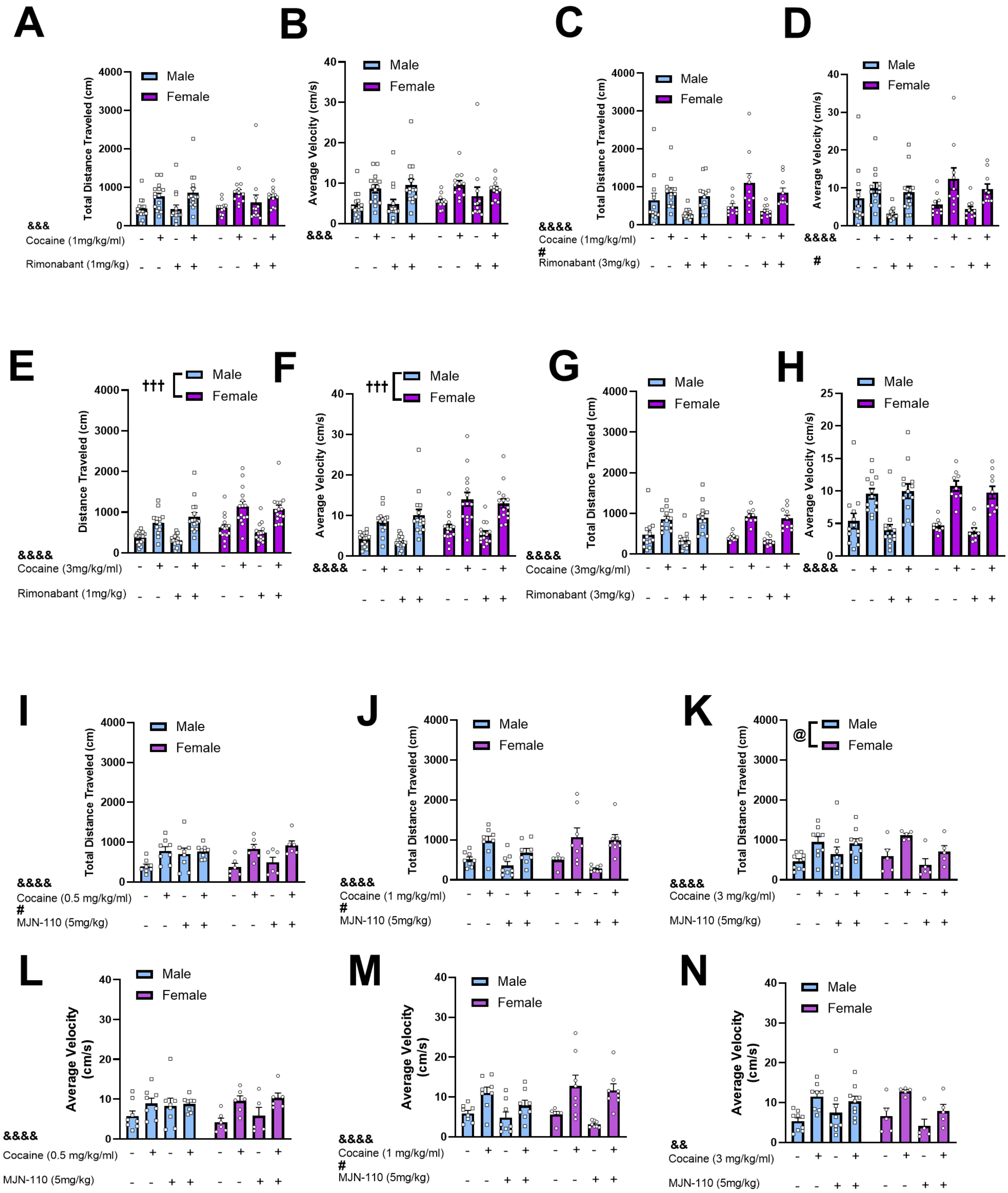

**Figure S1.** Locomotor activity is enhanced by cocaine treatment, and endocannabinoid modulation modestly reduces overall locomotor activity. (A) Cocaine (1mg/kg/ml) infusions significantly increased total distance traveled compared to saline but Rimonabant (1mg/kg) pretreatment had no significant effect on distance traveled or (B) average velocity. (C) A higher dose Rimonabant (3mg/kg) significantly reduced distance traveled compared to vehicle but did not attenuate Cocaine (1mg/kg/ml)-induced increases in distance traveled. (D) A higher dose Rimonabant (3mg/kg) significantly reduced average velocity compared to vehicle but did not attenuate Cocaine (1mg/kg/ml)-induced increases in average velocity. (E) Cocaine infusions (3mg/kg/ml) significantly increased distance traveled and average velocity (F) compared to saline, and females overall showed greater distance traveled and average velocity. No effects were observed from pretreatment with Rimonabant (1mg/kg) compared to vehicle. (G) Cocaine infusions (3mg/kg/ml) significantly increased distance traveled and average velocity (H) compared to saline, but Rimonabant (3mg/kg) did not significantly affect distance traveled or average velocity. No sex effects were observed for this testing condition. (I) MJN-110 significantly increased distance traveled compared to vehicle but did not impact cocaine (0.5 mg/kg/ml)-induced increases in distance traveled. (J) MJN-110 significantly decreased distance traveled compared to vehicle but did not impact cocaine (1 mg/kg/ml)-induced increases in distance traveled. (K) Cocaine (3mg/kg/ml) increased distance traveled and there were sex-specific impacts of MJN-110 on distance traveled compared to vehicle. However, MJN-110 did not impact cocaine-induced increases in distance traveled (L) Cocaine (0.5mg/kg/ml) increased average velocity and pretreatment with MJN-110 (5mg/kg) had no effect on average velocity. (M) MJN-110 (5mg/kg) pretreatment decreased average velocity compared to vehicle but had no effect on cocaine (1 mg/kg/ml)-induced increases in average velocity. (N) Cocaine (3mg/kg/ml) increased average velocity but pretreatment with MJN-110 (5mg/kg) had no effect. Data are presented as mean ± SEM; &&$p < .01$, &&$p < .001$, &&&&$p < .0001$ for Cocaine compared to saline across sexes; #$p < .05$ for Rimonabant or MJN-110 compared to vehicle across sexes; †††$p < .001$ for main effect between males and females.

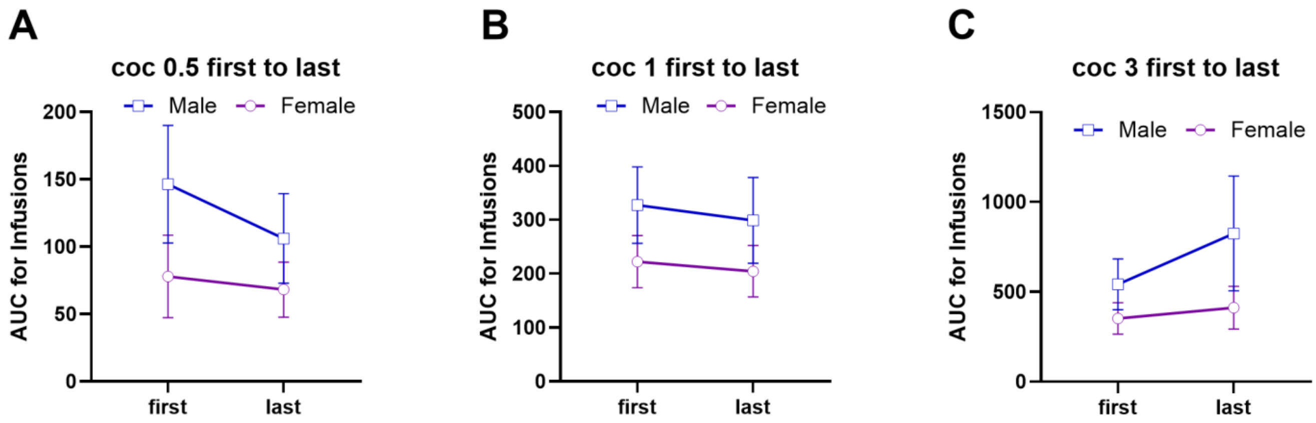

**Figure S2**. The cumulative effects of repeated cocaine treatment across days does not significantly affect the overall DA response to cocaine infusions. (A) Males (n = 7) and females (n = 5) do not show any sex effects or effects of cumulative cocaine exposure on the DA response to a low dose cocaine (0.5mg/kg/ml) infusion. (B) Males (n = 6) and females (n = 5) do not show any sex effects or effects of cumulative cocaine exposure on the DA response to a moderate dose cocaine (1mg/kg/ml) infusion. (C) Males (n = 8) and females (n = 6) do not show any sex effects or effects of cumulative cocaine exposure on the DA response to a high dose cocaine (3mg/kg/ml) infusion. Data are presented as mean ± SEM.

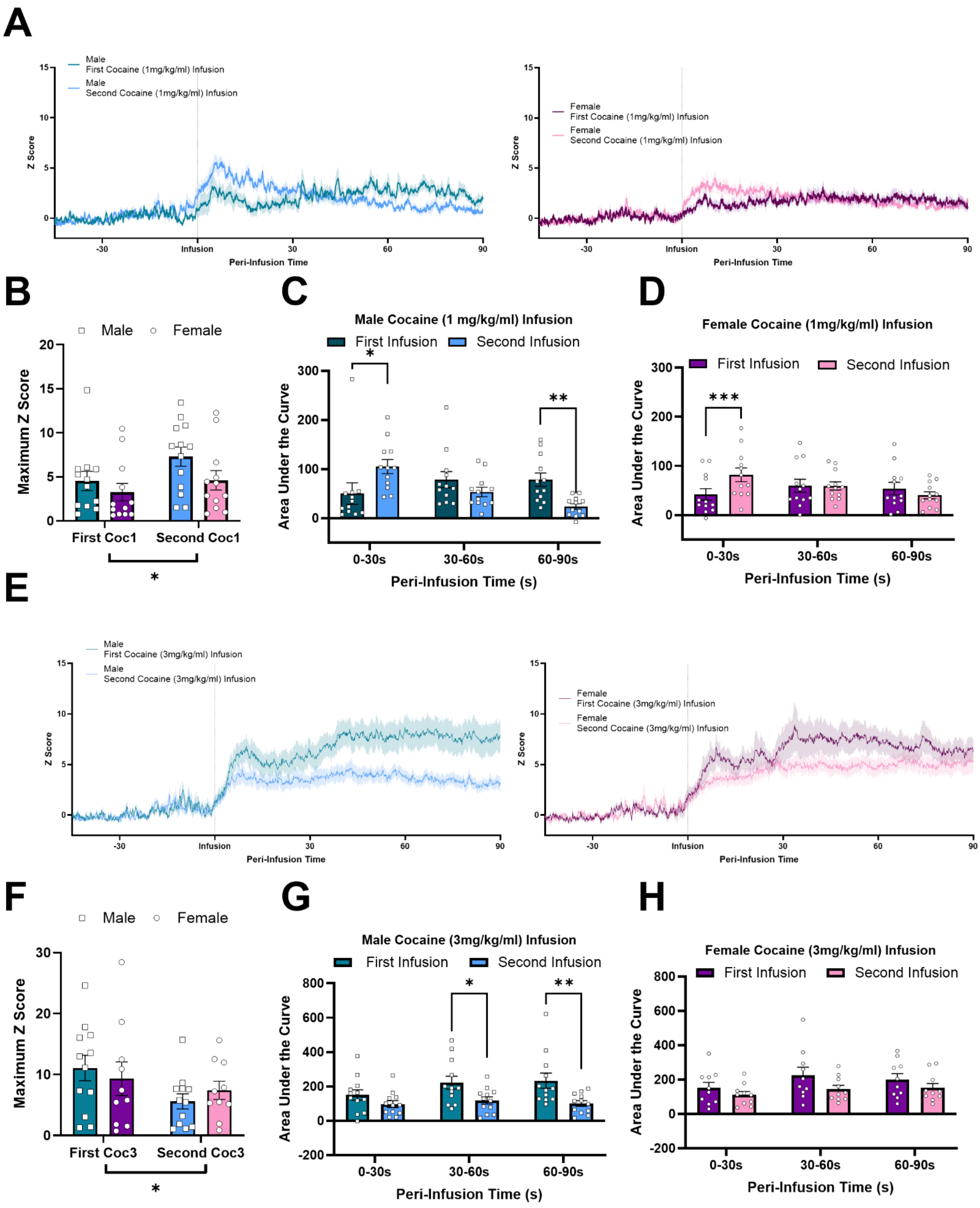
A
Male First Cocaine (1mg/kg/ml) Infusion
Male Second Cocaine (1mg/kg/ml) Infusion
Female First Cocaine (1mg/kg/ml) Infusion
Female Second Cocaine (1mg/kg/ml) Infusion
Z Score
Peri-Infusion Time
B
Male
Female
Maximum Z Score
First Coc1
Second Coc1
C
Male Cocaine (1 mg/kg/ml) Infusion
First Infusion
Second Infusion
Area Under the Curve
0-30s
30-60s
60-90s
Peri-Infusion Time (s)
D
Female Cocaine (1mg/kg/ml) Infusion
E
Male First Cocaine (3mg/kg/ml) Infusion
Male Second Cocaine (3mg/kg/ml) Infusion
Female First Cocaine (3mg/kg/ml) Infusion
Female Second Cocaine (3mg/kg/ml) Infusion
F
First Coc3
Second Coc3
G
Male Cocaine (3mg/kg/ml) Infusion
H
Female Cocaine (3mg/kg/ml) Infusion

**Figure S3.** Repeated cocaine infusions differentially change the temporal dynamics and maximum amplitude of $NAc_{ms}$ DA signaling depending on cocaine dose. (A) Average Z-scored traces of male (n = 13, left) and female (n = 12, right) given repeated infusions of a moderate dose of cocaine (1mg/kg/ml, i.v.). (B) Across sexes, a second infusion of a moderate dose of cocaine showed an enhanced Maximum Z score when compared to the first infusion. (C) Area under the curve (AUC) analysis of moderate dose cocaine response over time for males. The second infusion showed an initial enhancement followed by a more rapid decay of signal. (D) AUC analysis of moderate dose cocaine response over time for females. The second infusion showed an initial enhancement of signal. (E) Average Z-scored traces of male (n = 12, left) and female (n = 10, right) given repeated infusions of a high dose of cocaine (3mg/kg/ml, i.v.). (F) Across sexes, a second infusion of a high dose of cocaine showed reduced maximum Z score when compared to the first infusion. (G) AUC analysis of a high dose cocaine response over time for males. The second infusion showed reduced cocaine response for the later time bins. (H) AUC analysis of a high dose cocaine response over time for females. There were no significant differences between first and second infusion across the time bins. Data are presented as mean ± SEM; *p < .05, **p < .01, ***p < .001 for first infusion compared to second infusion.

### Tables

| *Comparison* | *Statistical Test* | *Statistic* | *p-value* |
|---|---|---|---|
| **B** | | | |
| **Area Under the Curve Dose Response to Cocaine** | Two-way RM ANOVA | | |
| Cocaine doses (within) x Sex (between) | Interaction | F (3, 51) = 2.46 | P=0.073 |
| Cocaine dose | Main effect | F (3, 51) = 43.51 | **P<0.000** |
| Sex: male, n = 10; female, n = 9 | Main effect | F (1, 17) = 0.49 | P=0.492 |
| Saline vs. Cocaine (0.5mg/kg) | Holm-Šídák | | **P=0.024** |
| Saline vs. Cocaine (1mg/kg) | Holm-Šídák | | **P<0.001** |
| Saline vs. Cocaine (3mg/kg) | Holm-Šídák | | **P<0.001** |
| Cocaine (0.5mg/kg) vs Cocaine (1mg/kg) | Holm-Šídák | | **P=0.024** |
| Cocaine (0.5mg/kg) vs Cocaine (3mg/kg) | Holm-Šídák | | **P<0.001** |
| Cocaine (1mg/kg) vs Cocaine (3mg/kg) | Holm-Šídák | | **P<0.001** |
| **C** | | | |
| **Distance Traveled Dose Response to Cocaine** | One-way RM ANOVA | | |
| Cocaine Dose: n = 7 | Main effect | F (3, 18) = 4.73 | **P=0.013** |
| Saline vs. Cocaine (0.5mg/kg) | Dunnett | | **P=0.017** |
| Saline vs. Cocaine (1mg/kg) | Dunnett | | **P=0.017** |
| Saline vs. Cocaine (3mg/kg) | Dunnett | | **P=0.017** |
| **D** | | | |
| **Average Velocity Dose Response to Cocaine** | One-way RM ANOVA | | |
| Cocaine Dose: n = 7 | Main effect | F (3, 18) = 4.53 | **P=0.016** |
| Saline vs. Cocaine (0.5mg/kg) | Dunnett | | P=0.125 |
| Saline vs. Cocaine (1mg/kg) | Dunnett | | **P=0.047** |
| Saline vs. Cocaine (3mg/kg) | Dunnett | | **P=0.040** |

**Table S1.** Detailed statistics for Figure 2. Significant p-values are bolded.

| *Comparison* | *Statistical Test* | *Statistic* | *p-value* |
|---|---|---|---|
| **B** | | | |
| **Frequency Response to Rimonabant (1mg/kg)** | Two-way RM ANOVA | | |
| Drug Pretreatment (within) x Sex (between) | Interaction | F (1, 20) = 1.38 | P=0.255 |
| Drug (Rimonabant 1mg/kg) | Main effect | F (1, 20) = 0.56 | P=0.464 |
| Sex: male, n = 12; female, n = 11 | Main effect | F (1, 20) = 0.25 | P=0.621 |
| **C** | | | |
| **Average Amplitude for Rimonabant (1mg/kg)** | Two-way RM ANOVA | | |
| Drug Pretreatment (within) x Sex (between) | Interaction | F (1, 19) = 3.70 | P=0.070 |
| Drug (Rimonabant 1mg/kg) | Main effect | F (1, 19) = 1.21 | P=0.286 |
| Sex: male, n = 12; female, n = 11 | Main effect | F (1, 19) = 0.01 | P=0.915 |
| **E** | | | |
| **Frequency Response to Rimonabant (3mg/kg)** | Two-way RM ANOVA | | |
| Drug Pretreatment (within) x Sex (between) | Interaction | F (1, 24) = 0.01 | P=0.972 |
| Drug (Rimonabant 3mg/kg) | Main effect | F (1, 24) = 0.25 | P=0.623 |
| Sex: male, n = 16; female, n = 12 | Main effect | F (1, 24) = 0.01 | P=0.931 |
| **F** | | | |
| **Average Amplitude for Rimonabant (3mg/kg)** | Two-way RM ANOVA | | |
| Drug Pretreatment (within) x Sex (between) | Interaction | F (1, 22) = 0.37 | P=0.549 |
| Drug (Rimonabant 3mg/kg) | Main effect | F (1, 22) = 1.75 | P=0.200 |
| Sex: male, n = 16; female, n = 12 | Main effect | F (1, 22) = 0.04 | P=0.853 |
| **H** | | | |
| **Frequency Response to MJN-110 (5mg/kg)** | Two-way RM ANOVA | | |
| Drug Pretreatment (within) x Sex (between) | Interaction | F (1, 13) = 0.02 | P=0.884 |
| Drug (MJN-110 5mg/kg) | Main effect | F (1, 13) = 0.03 | P=0.873 |
| Sex: male, n = 8; female, n = 8 | Main effect | F (1, 13) = 0.06 | P=0.811 |
| **I** | | | |
| **Average Amplitude for MJN-110 (5mg/kg)** | Two-way RM ANOVA | | |
| Drug Pretreatment (within) x Sex (between) | Interaction | F (1, 13) = 2.03 | P=0.178 |
| Drug (MJN-110 5mg/kg) | Main effect | F (1, 13) = 3.27 | P=0.094 |
| Sex: male, n = 7; female, n = 8 | Main effect | F (1, 13) = 2.27 | P=0.156 |

**Table S2.** Detailed statistics for Figure 3. Significant p-values are bolded.

| *Comparison* | *Statistical Test* | *Statistic* | *p-value* |
|---|---|---|---|
| **C-D (left)** | | | |
| **Area Under the Curve of Cocaine (1mg/kg) + Rimonabant (1mg/kg)** | Three-way RM ANOVA | | |
| Time (within) x Sex (between) x Drug (within) | Interaction | F (2, 28) = 0.68 | P=0.516 |
| Time x Sex | Interaction | F (2, 28) = 0.88 | P=0.427 |
| Time x Drug | Interaction | F (2, 28) = 0.26 | P=0.769 |
| Sex x Drug | Interaction | F (1, 14) = 0.01 | P=0.937 |
| Time | Main effect | F (2, 28) = 0.60 | P=0.555 |
| Sex: male, n = 8; female, n = 8 | Main effect | F (1, 14) = 0.38 | P=0.550 |
| Drug | Main effect | F (1, 14) = 0.89 | P=0.363 |
| **E (left)** | | | |
| **Maximum Amplitude of Cocaine (1mg/kg) Response** | Two-way RM ANOVA | | |
| Sex (between) x Drug (within) | Interaction | F (1, 15) = 4.90 | **P=0.043** |
| Sex: male, n = 8; female, n = 8 | Main effect | F (1, 15) = 0.17 | P=0.690 |
| Drug | Main effect | F (1, 15) = 0.29 | P=0.599 |
| Drug effect: male | Holm-Šídák | | P=0.241 |
| Drug effect: female | Holm-Šídák | | P=0.151 |

**Table S3.** Detailed statistics for Figure 4 (left). Significant p-values are bolded.

| *Comparison* | *Statistical Test* | *Statistic* | *p-value* |
|---|---|---|---|
| **C-D (right)** | | | |
| **Area Under the Curve of Cocaine (1mg/kg) + Rimonabant (3mg/kg)** | Three-way RM ANOVA | | |
| Time (within) x Sex (between) x Drug (within) | Interaction | F (2, 30) = 0.96 | P=0.396 |
| Time x Sex | Interaction | F (2, 30) = 2.49 | P=0.100 |
| Time x Drug | Interaction | F (2, 30) = 1.46 | P=0.249 |
| Sex x Drug | Interaction | F (1, 15) = 0.34 | P=0.570 |
| Time | Main effect | F (2, 30) = 0.11 | P=0.896 |
| Sex: male, n = 8; female, n = 9 | Main effect | F (1, 15) = 0.51 | P=0.484 |
| Drug | Main effect | F (1, 15) = 6.90 | **P=0.019** |
| *Comparison* | *Statistical Test* | *Statistic* | *p-value* |
| **C** | | | |
| **Male Area Under the Curve of Cocaine (1mg/kg) + Rimonabant (3mg/kg)** | Two-way RM ANOVA | | |
| Time (within) x Drug (within): n = 8 | Interaction | F (2, 14) = 0.04 | P=0.965 |
| Time | Main effect | F (2, 14) = 2.25 | P=0.142 |
| Drug | Main effect | F (1, 7) = 1.21 | P=0.308 |
| *Comparison* | *Statistical Test* | *Statistic* | *p-value* |
| **D** | | | |
| **Female Area Under the Curve of Cocaine (1mg/kg) + Rimonabant (3mg/kg)** | Two-way RM ANOVA | | |
| Time (within) x Drug (within): n = 9 | Interaction | F (2, 16) = 2.02 | P=0.166 |
| Time | Main effect | F (2, 16) = 0.79 | P=0.473 |
| Drug | Main effect | F (1, 8) = 12.22 | **P=0.008** |
| *Comparison* | *Statistical Test* | *Statistic* | *p-value* |
| **E** | | | |
| **Maximum Amplitude of Cocaine (3mg/kg) Response** | Two-way RM ANOVA | | |
| Sex (between) x Drug (within) | Interaction | F (1, 14) = 2.26 | P=0.155 |
| Sex: male, n = 8; female, n = 9 | Main effect | F (1, 14) = 0.01 | P=0.928 |
| Drug | Main effect | F (1, 14) = 5.65 | **P=0.032** |
| Drug effect: male | Holm-Šídák | | P=0.570 |
| Drug effect: female | Holm-Šídák | | **P=0.022** |

**Table S4.** Detailed statistics for Figure 4 (right). Significant p-values are bolded.

| *Comparison* | *Statistical Test* | *Statistic* | *p-value* |
|---|---|---|---|
| **C-D (left)** | | | |
| **Area Under the Curve of Cocaine (3mg/kg) + Rimonabant (1mg/kg)** | Three-way RM ANOVA | | |
| Time (within) x Sex (between) x Drug (within) | Interaction | F (2, 42) = 2.99 | P=0.061 |
| Time x Sex | Interaction | F (2, 42) = 3.04 | P=0.059 |
| Time x Drug | Interaction | F (2, 42) = 2.18 | P=0.125 |
| Sex x Drug | Interaction | F (1, 21) = 4.87 | **P=0.039** |
| Time | Main effect | F (2, 42) = 8.91 | **P=0.001** |
| Sex: male, n = 11; female, n = 12 | Main effect | F (1, 21) = 0.14 | P=0.708 |
| Drug | Main effect | F (1, 21) = 6.82 | **P=0.016** |
| *Comparison* | *Statistical Test* | *Statistic* | *p-value* |
| **C (left)** | | | |
| **Male Area Under the Curve of Cocaine (3mg/kg) + Rimonabant (1mg/kg)** | Two-way RM ANOVA | | |
| Time (within) x Drug (within): n = 11 | Interaction | F (2, 20) = 0.12 | P=0.892 |
| Time | Main effect | F (2, 20) = 2.54 | P=0.104 |
| Drug | Main effect | F (1, 10) = 0.06 | P=0.813 |
| *Comparison* | *Statistical Test* | *Statistic* | *p-value* |
| **D (left)** | | | |
| **Female Area Under the Curve of Cocaine (3mg/kg) + Rimonabant (1mg/kg)** | Two-way RM ANOVA | | |
| Time (within) x Drug (within): n = 12 | Interaction | F (2, 22) = 3.42 | **P=0.051** |
| Time | Main effect | F (2, 22) = 7.28 | **P=0.004** |
| Drug | Main effect | F (1, 11) = 17.59 | **P=0.002** |
| *Comparison* | *Statistical Test* | *Statistic* | *p-value* |
| **E (left)** | | | |
| **Maximum Amplitude of Cocaine (3mg/kg) Response** | Two-way RM ANOVA | | |
| Sex (between) x Drug (within) | Interaction | F (1, 22) = 14.34 | **P=0.001** |
| Sex: male, n = 11; female, n = 12 | Main effect | F (1, 22) = 0.57 | P=0.459 |
| Drug | Main effect | F (1, 22) = 34.41 | **P<0.001** |
| Drug effect: male | Holm-Šídák's multiple comparisons test | | P=0.156 |
| Drug effect: female | Holm-Šídák's multiple comparisons test | | **P<0.001** |

**Table S5.** Detailed statistics for Figure 5 (left). Significant p-values are bolded.

| *Comparison* | *Statistical Test* | *Statistic* | *p-value* |
|---|---|---|---|
| **C-D (right)** | | | |
| **Area Under the Curve of Cocaine (3mg/kg) + Rimonabant (3mg/kg)** | Three-way RM ANOVA | | |
| Time (within) x Sex (between) x Drug (within) | Interaction | F (2, 24) = 0.65 | P=0.532 |
| Time x Sex | Interaction | F (2, 24) = 0.33 | P=0.721 |
| Time x Drug | Interaction | F (2, 24) = 0.53 | P=0.595 |
| Sex x Drug | Interaction | F (1, 12) = 2.87 | P=0.116 |
| Time | Main effect | F (2, 24) = 0.88 | P=0.428 |
| Sex: male, n = 7; female, n = 7 | Main effect | F (1, 12) = 4.67 | **P=0.052** |
| Drug | Main effect | F (1, 12) = 31.14 | **P<0.001** |
| **C (right)** | | | |
| **Male Area Under the Curve of Cocaine (3mg/kg) + Rimonabant (3m** | Two-way RM ANOVA | | |
| Time (within) x Drug (within): n = 7 | Interaction | F (2, 12) = 0.29 | P=0.753 |
| Time | Main effect | F (2, 12) = 1.18 | P=0.341 |
| Drug | Main effect | F (1, 6) = 17.86 | **P=0.006** |
| **D (right)** | | | |
| **Female Area Under the Curve of Cocaine (3mg/kg) + Rimonabant (** | Two-way RM ANOVA | | |
| Time (within) x Drug (within): n = 7 | Interaction | F (2, 12) = 0.92 | P=0.424 |
| Time | Main effect | F (2, 12) = 0.06 | P=0.938 |
| Drug | Main effect | F (1, 6) = 14.57 | **P=0.009** |
| **E (right)** | | | |
| Maximum Amplitude of Cocaine (3mg/kg) Response | Two-way RM ANOVA | | |
| Sex (between) x Drug (within) | Interaction | F (1, 12) = 0.74 | P=0.406 |
| Sex: male, n = 7; female, n = 7 | Main effect | F (1, 12) = 1.33 | P=0.271 |
| Drug | Main effect | F (1, 12) = 22.15 | **P=0.001** |
| Drug effect: male | Holm-Šídák's multiple comparisons test | | **P=0.004** |
| Drug effect: female | Holm-Šídák's multiple comparisons test | | **P=0.019** |

**Table S6.** Detailed statistics for Figure 5 (right). Significant p-values are bolded.

| *Comparison* | *Statistical Test* | *Statistic* | *p-value* |
|---|---|---|---|
| **C-D** | | | |
| **Area Under the Curve of Cocaine (0.5mg/kg) + MJN-110 (5mg/kg)** | Three-way RM ANOVA | | |
| Time (within) x Sex (between) x Drug (within): male, n = 8; female, n = 6 | Interaction | F (2, 24) = 6.77 | **P=0.005** |
| Time x Sex | Interaction | F (2, 24) = 0.57 | P=0.573 |
| Time x Drug | Interaction | F (2, 24) = 1.28 | P=0.297 |
| Sex x Drug | Interaction | F (1, 12) = 0.61 | P=0.449 |
| Time | Main effect | F (2, 24) = 1.75 | P=0.195 |
| Sex | Main effect | F (1, 12) = 0.16 | P=0.695 |
| Drug | Main effect | F (1, 12) = 3.42 | P=0.089 |
| **C** | | | |
| **Male Area Under the Curve of Cocaine (0.5mg/kg) + MJN-110 (5mg/kg)** | Two-way RM ANOVA | | |
| Time (within) x Drug (within): n = 8 | Interaction | F (2, 14) = 7.83 | **P=0.005** |
| Time | Main effect | F (2, 14) = 0.49 | P=0.622 |
| Drug | Main effect | F (1, 7) = 0.46 | P=0.518 |
| **D** | | | |
| **Female Area Under the Curve of Cocaine (0.5mg/kg) + MJN-110 (5mg/kg)** | Two-way RM ANOVA | | |
| Time (within) x Drug (within): n = 6 | Interaction | F (2, 10) = 2.13 | P=0.169 |
| Time | Main effect | F (2, 10) = 1.13 | P=0.361 |
| Drug | Main effect | F (1, 5) = 7.55 | **P=0.041** |
| **E** | | | |
| **Maximum Amplitude of Cocaine (0.5mg/kg) + MJN-110 (5mg/kg) Response** | Two-way RM ANOVA | | |
| Sex (between) x Drug (within): male, n = 8; female, n = 6 | Interaction | F (1, 13) = 0.96 | P=0.346 |
| Sex | Main effect | F (1, 13) = 0.13 | P=0.720 |
| Drug | Main effect | F (1, 13) = 2.08 | P=0.173 |

**Table S7.** Detailed statistics for Figure 6. Significant p-values are bolded.

| ***Comparison*** | ***Statistical Test*** | ***Statistic*** | ***p-value*** |
|---|---|---|---|
| **C-D** | | | |
| **Area Under the Curve of Cocaine (1mg/kg) + MJN-110 (5mg/kg)** | Three-way RM ANOVA | | |
| Time (within) x Sex (between) x Drug (within): male, n = 7; female, n = 8 | Interaction | F (2, 26) = 0.30 | P=0.743 |
| Time x Sex | Interaction | F (2, 26) = 0.55 | P=0.584 |
| Time x Drug | Interaction | F (2, 26) = 0.82 | P=0.453 |
| Sex x Drug | Interaction | F (1, 13) = 0.07 | P=0.799 |
| Time | Main effect | F (2, 26) = 0.25 | P=0.781 |
| Sex | Main effect | F (1, 13) = 0.11 | P=0.751 |
| Drug | Main effect | F (1, 13) = 8.87 | **P=0.011** |
| **C** | | | |
| **Male Area Under the Curve of Cocaine (1mg/kg) + MJN-110 (5mg/kg)** | Two-way RM ANOVA | | |
| Time (within) x Drug (within): n = 7 | Interaction | F (2, 12) = 0.37 | P=0.697 |
| Time | Main effect | F (2, 12) = 0.71 | P=0.514 |
| Drug | Main effect | F (1, 6) = 3.01 | P=0.134 |
| **D** | | | |
| **Female Area Under the Curve of Cocaine (1mg/kg) + MJN-110 (5mg/kg)** | Two-way RM ANOVA | | |
| Time (within) x Drug (within): n = 8 | Interaction | F (2, 14) = 0.80 | P=0.470 |
| Time | Main effect | F (2, 14) = 0.07 | P=0.935 |
| Drug | Main effect | F (1, 7) = 6.46 | **P=0.039** |
| **E** | | | |
| **Maximum Amplitude of Cocaine (1mg/kg) + MJN-110 (5mg/kg) Response** | Two-way RM ANOVA | | |
| Sex (between) x Drug (within): male, n = 7; female, n = 8 | Interaction | F (1, 13) = 0.19 | P=0.666 |
| Sex | Main effect | F (1, 13) = 0.17 | P=0.687 |
| Drug | Main effect | F (1, 13) = 5.25 | **P=0.039** |
| Drug effect: male | Holm-Šídák | | P=0.161 |
| Drug effect: female | Holm-Šídák | | P=0.358 |

**Table S8.** Detailed statistics for Figure 7. Significant p-values are bolded.

| ***Comparison*** | ***Statistical Test*** | ***Statistic*** | ***p-value*** |
|---|---|---|---|
| **C-D** | | | |
| **Area Under the Curve of Cocaine (3mg/kg) + MJN-110 (5mg/kg)** | Three-way RM ANOVA | | |
| Time (within) x Sex (between) x Drug (within): male, n = 8; female, n = 8 | Interaction | F (2, 28) = 0.05 | P=0.951 |
| Time x Sex | Interaction | F (2, 28) = 0.45 | P=0.641 |
| Time x Drug | Interaction | F (2, 28) = 0.39 | P=0.682 |
| Sex x Drug | Interaction | F (1, 14) = 7.45 | **P=0.016** |
| Time | Main effect | F (2, 28) = 2.49 | P=0.101 |
| Sex | Main effect | F (1, 14) = 0.11 | P=0.744 |
| Drug | Main effect | F (1, 14) = 1.52 | P=0.239 |
| **C** | | | |
| **Male Area Under the Curve of Cocaine (3mg/kg) + MJN-110 (5mg/kg)** | Two-way RM ANOVA | | |
| Time (within) x Drug (within): n = 8 | Interaction | F (2, 14) = 0.13 | P=0.875 |
| Time | Main effect | F (2, 14) = 0.89 | P=0.431 |
| Drug | Main effect | F (1, 7) = 1.52 | P=0.258 |
| **D** | | | |
| **Female Area Under the Curve of Cocaine (3mg/kg) + MJN-110 (5mg/kg)** | Two-way RM ANOVA | | |
| Time (within) x Drug (within): n = 8 | Interaction | F (2, 14) = 0.28 | P=0.761 |
| Time | Main effect | F (2, 14) = 2.18 | P=0.150 |
| Drug | Main effect | F (1, 7) = 6.22 | **P=0.041** |
| **E** | | | |
| **Maximum Amplitude of Cocaine (3mg/kg) + MJN-110 (5mg/kg) Response** | Two-way RM ANOVA | | |
| Sex (between) x Drug (within): male, n = 8; female, n = 8 | Interaction | F (1, 13) = 8.23 | **P=0.013** |
| Sex | Main effect | F (1, 13) = 0.13 | P=0.729 |
| Drug | Main effect | F (1, 13) = 0.24 | P=0.632 |
| Drug effect: male | Holm-Šídák | | P=0.128 |
| Drug effect: female | Holm-Šídák | | P=0.057 |

**Table S9.** Detailed statistics for Figure 8. Significant p-values are bolded.

| ***Comparison*** | ***Statistical Test*** | ***Statistic*** | ***p-value*** |
|---|---|---|---|
| **A** | | | |
| Estrus Effect on Cocaine Dose Response | Two-way ANOVA | | |
| Cocaine Dose (between) x Estrus (between) | Interaction | F (2, 53) = 4.20 | **P=0.020** |
| Cocaine Dose | Main effect | F (2, 53) = 11.57 | **P<0.001** |
| Estrus | Main effect | F (1, 53) = 0.82 | P=0.370 |
| Non-Estrus vs Estrus, Cocaine (0.5mg/kg/ml): non-estrus, n = 6; estrus, n = 5 | Holm-Šídák | | P=0.905 |
| Non-Estrus vs Estrus, Cocaine (1mg/kg/ml): non-estrus, n = 18; estrus, n = 7 | Holm-Šídák | | P=0.520 |
| Non-Estrus vs Estrus, Cocaine (3mg/kg/ml): non-estrus, n = 16; estrus, n = 7 | Holm-Šídák | | **P=0.015** |

**Table S10.** Detailed statistics for Figure 9. Significant p-values are bolded.

| *Comparison* | *Statistical Test* | *Statistic* | *p-value* |
|---|---|---|---|
| **A** | | | |
| **Total Distance Traveled: Cocaine (1mg/kg/ml) + Rimonabant (1mg/kg)** | Three-way RM ANOVA | | |
| Cocaine (within) x Sex (between) x Drug (within) | Interaction | F (1, 25) = 2.88 | P=0.102 |
| Cocaine x Sex | Interaction | F (1, 25) = 0.52 | P=0.478 |
| Cocaine x Drug | Interaction | F (1, 25) = 0.50 | P=0.487 |
| Sex x Drug | Interaction | F (1, 25) = 0.04 | P=0.845 |
| Cocaine | Main effect | F (1, 25) = 19.48 | **P<0.001** |
| Sex: male, n = 15; female, n = 12 | Main effect | F (1, 25) = 0.41 | P=0.526 |
| Drug | Main effect | F (1, 25) = 0.11 | P=0.745 |
| **B** | | | |
| **Average Velocity: Cocaine (1mg/kg/ml) + Rimonabant (1mg/kg)** | Three-way RM ANOVA | | |
| Cocaine (within) x Sex (between) x Drug (within) | Interaction | F (1, 25) = 2.00 | P=0.171 |
| Cocaine x Sex | Interaction | F (1, 25) = 0.82 | P=0.374 |
| Cocaine x Drug | Interaction | F (1, 25) = 0.53 | P=0.471 |
| Sex x Drug | Interaction | F (1, 25) = 0.09 | P=0.763 |
| Cocaine | Main effect | F (1, 25) = 20.48 | **P<0.001** |
| Sex: male, n = 15; female, n = 12 | Main effect | F (1, 25) = 0.69 | P=0.415 |
| Drug | Main effect | F (1, 25) = 0.09 | P=0.772 |
| **C** | | | |
| **Total Distance Traveled: Cocaine (1mg/kg/ml) + Rimonabant (3mg/kg)** | Three-way RM ANOVA | | |
| Cocaine (within) x Sex (between) x Drug (within) | Interaction | F (1, 21) = 1.23 | P=0.281 |
| Cocaine x Sex | Interaction | F (1, 21) = 1.40 | P=0.250 |
| Cocaine x Drug | Interaction | F (1, 21) = 0.19 | P=0.734 |
| Sex x Drug | Interaction | F (1, 21) = 0.09 | P=0.764 |
| Cocaine | Main effect | F (1, 21) = 24.29 | **P<0.001** |
| Sex: male, n = 12; female, n = 10 | Main effect | F (1, 21) = 0.38 | P=0.543 |
| Drug | Main effect | F (1, 21) = 5.42 | **P=0.030** |
| **D** | | | |
| **Average Velocity: Cocaine (1mg/kg/ml) + Rimonabant (3mg/kg)** | Three-way RM ANOVA | | |
| Cocaine (within) x Sex (between) x Drug (within) | Interaction | F (1, 21) = 1.20 | P=0.285 |
| Cocaine x Sex | Interaction | F (1, 21) = 0.81 | P=0.378 |
| Cocaine x Drug | Interaction | F (1, 21) = 0.17 | P=0.683 |
| Sex x Drug | Interaction | F (1, 21) = 0.10 | P=0.753 |
| Cocaine | Main effect | F (1, 21) = 23.15 | **P<0.001** |
| Sex: male, n = 12; female, n = 10 | Main effect | F (1, 21) = 0.30 | P=0.588 |
| Drug | Main effect | F (1, 21) = 4.67 | **P=0.042** |
| **E** | | | |
| **Total Distance Traveled: Cocaine (3mg/kg/ml) + Rimonabant (1mg/kg)** | Three-way RM ANOVA | | |
| Cocaine (within) x Sex (between) x Drug (within) | Interaction | F (1, 26) = 0.11 | P=0.747 |
| Cocaine x Sex | Interaction | F (1, 26) = 1.65 | P=0.211 |
| Cocaine x Drug | Interaction | F (1, 26) = 0.47 | P=0.499 |
| Sex x Drug | Interaction | F (1, 26) = 1.21 | P=0.281 |
| Cocaine | Main effect | F (1, 26) = 133.7 | **P<0.001** |
| Sex: male, n = 13; female, n = 15 | Main effect | F (1, 26) = 15.71 | **P<0.001** |
| Drug | Main effect | F (1, 26) = 0.9040 | P=0.353 |
| **F** | | | |
| **Average Velocity: Cocaine (3mg/kg/ml) + Rimonabant (1mg/kg)** | Three-way RM ANOVA | | |
| Cocaine (within) x Sex (between) x Drug (within) | Interaction | F (1, 26) = 0.14 | P=0.708 |
| Cocaine x Sex | Interaction | F (1, 26) = 2.31 | P=0.140 |
| Cocaine x Drug | Interaction | F (1, 26) = 0.32 | P=0.579 |
| Sex x Drug | Interaction | F (1, 26) = 1.17 | P=0.290 |
| Cocaine | Main effect | F (1, 26) = 88.69 | **P<0.001** |
| Sex: male, n = 13; female, n = 15 | Main effect | F (1, 26) = 16.04 | **P<0.001** |
| Drug | Main effect | F (1, 26) = 0.90 | P=0.351 |
| **G** | | | |
| **Total Distance Traveled: Cocaine (3mg/kg/ml) + Rimonabant (3mg/kg)** | Three-way RM ANOVA | | |
| Cocaine (within) x Sex (between) x Drug (within) | Interaction | F (1, 19) = 1.15 | P=0.296 |
| Cocaine x Sex | Interaction | F (1, 19) = 0.22 | P=0.645 |
| Cocaine x Drug | Interaction | F (1, 19) = 0.57 | P=0.461 |
| Sex x Drug | Interaction | F (1, 19) = 0.11 | P=0.747 |
| Cocaine | Main effect | F (1, 19) = 61.84 | **P<0.001** |
| Sex: male, n = 13; female, n = 8 | Main effect | F (1, 19) = 0.01 | P=0.905 |
| Drug | Main effect | F (1, 19) = 1.46 | P=0.242 |

| | | | |
|---|---|---|---|
| **H** | | | |
| **Average Velocity: Cocaine (3mg/kg/ml) + Rimonabant (3mg/kg)** | Three-way RM ANOVA | | |
| Cocaine (within) x Sex (between) x Drug (within) | Interaction | F (1, 19) = 1.49 | P=0.237 |
| Cocaine x Sex | Interaction | F (1, 19) = 0.36 | P=0.557 |
| Cocaine x Drug | Interaction | F (1, 19) = 0.19 | P=0.666 |
| Sex x Drug | Interaction | F (1, 19) = 0.19 | P=0.667 |
| Cocaine | Main effect | F (1, 19) = 61.63 | **P<0.001** |
| Sex: male, n = 13; female, n = 8 | Main effect | F (1, 19) = 0.00 | P=0.995 |
| Drug | Main effect | F (1, 19) = 1.393 | P=0.253 |
| **I** | | | |
| **Total Distance Traveled: Cocaine (0.5mg/kg/ml) +MJN-110 (5mg/kg)** | Three-way RM ANOVA | | |
| Cocaine (within) x Sex (between) x Drug (within) | Interaction | F (1, 12) = 0.63 | P=0.443 |
| Cocaine x Sex | Interaction | F (1, 12) = 4.04 | P=0.067 |
| Cocaine x Drug | Interaction | F (1, 12) = 0.98 | P=0.342 |
| Sex x Drug | Interaction | F (1, 12) = 0.19 | P=0.674 |
| Cocaine | Main effect | F (1, 12) = 38.88 | **P<0.001** |
| Sex: male, n = 8; female, n = 6 | Main effect | F (1, 12) = 0.00 | P=0.967 |
| Drug | Main effect | F (1, 12) = 6.79 | **P=0.023** |
| **J** | | | |
| **Total Distance Traveled: Cocaine (1mg/kg/ml) +MJN-110 (5mg/kg)** | Three-way RM ANOVA | | |
| Cocaine (within) x Sex (between) x Drug (within) | Interaction | F (1, 12) = 1.70 | P=0.217 |
| Cocaine x Sex | Interaction | F (1, 12) = 0.88 | P=0.367 |
| Cocaine x Drug | Interaction | F (1, 12) = 0.04 | P=0.852 |
| Sex x Drug | Interaction | F (1, 12) = 0.28 | P=0.607 |
| Cocaine | Main effect | F (1, 12) = 41.18 | **P<0.001** |
| Sex: male, n = 8; female, n = 6 | Main effect | F (1, 12) = 0.00 | P=0.960 |
| Drug | Main effect | F (1, 12) = 8.96 | **P=0.011** |
| **K** | | | |
| **Total Distance Traveled: Cocaine (3mg/kg/ml) +MJN-110 (5mg/kg)** | Three-way RM ANOVA | | |
| Cocaine (within) x Sex (between) x Drug (within) | Interaction | F (1, 12) = 0.01 | P=0.937 |
| Cocaine x Sex | Interaction | F (1, 12) = 0.10 | P=0.753 |
| Cocaine x Drug | Interaction | F (1, 12) = 1.57 | P=0.234 |
| Sex x Drug | Interaction | F (1, 12) = 4.83 | **P=0.048** |
| Cocaine | Main effect | F (1, 12) = 23.80 | **P<0.001** |
| Sex: male, n = 9; female, n = 5 | Main effect | F (1, 12) = 0.09 | P=0.767 |
| Drug | Main effect | F (1, 12) = 1.87 | P=0.197 |
| **L** | | | |
| **Average Velocity: Cocaine (0.5mg/kg/ml) +MJN-110 (5mg/kg)** | Three-way RM ANOVA | | |
| Cocaine (within) x Sex (between) x Drug (within) | Interaction | F (1, 11) = 0.02 | P=0.897 |
| Cocaine x Sex | Interaction | F (1, 11) = 6.25 | **P=0.030** |
| Cocaine x Drug | Interaction | F (1, 11) = 0.72 | P=0.413 |
| Sex x Drug | Interaction | F (1, 11) = 0.24 | P=0.632 |
| Cocaine | Main effect | F (1, 11) = 37.46 | **P<0.001** |
| Sex: male, n = 8; female, n = 6 | Main effect | F (1, 11) = 0.27 | P=0.611 |
| Drug | Main effect | F (1, 11) = 2.86 | P=0.119 |
| **M** | | | |
| **Average Velocity: Cocaine (1mg/kg/ml) +MJN-110 (5mg/kg)** | Three-way RM ANOVA | | |
| Cocaine (within) x Sex (between) x Drug (within) | Interaction | F (1, 13) = 3.38 | P=0.089 |
| Cocaine x Sex | Interaction | F (1, 13) = 0.91 | P=0.358 |
| Cocaine x Drug | Interaction | F (1, 13) = 0.67 | P=0.428 |
| Sex x Drug | Interaction | F (1, 13) = 0.57 | P=0.462 |
| Cocaine | Main effect | F (1, 13) = 43.57 | **P<0.001** |
| Sex: male, n = 8; female, n = 6 | Main effect | F (1, 13) = 0.61 | P=0.449 |
| Drug | Main effect | F (1, 13) = 5.71 | **P=0.033** |
| **N** | | | |
| **Average Velocity: Cocaine (3mg/kg/ml) +MJN-110 (5mg/kg)** | Three-way RM ANOVA | | |
| Cocaine (within) x Sex (between) x Drug (within) | Interaction | F (1, 11) = 0.08 | P=0.786 |
| Cocaine x Sex | Interaction | F (1, 11) = 0.19 | P=0.669 |
| Cocaine x Drug | Interaction | F (1, 11) = 2.77 | P=0.124 |
| Sex x Drug | Interaction | F (1, 11) = 4.74 | P=0.052 |
| Cocaine | Main effect | F (1, 11) = 17.77 | **P=0.001** |
| Sex: male, n = 9; female, n = 5 | Main effect | F (1, 11) = 0.51 | P=0.491 |
| Drug | Main effect | F (1, 11) = 2.28 | P=0.159 |

**Table S11.** Detailed statistics for Figure S1. Significant p-values are bolded.

| *Comparison* | *Statistical Test* | *Statistic* | *p-value* |
|---|---|---|---|
| **A** | | | |
| **AUC for coc (0.5mg/kg/ml) infusions from first to last** | Two-way RM ANOVA | | |
| Sex (between) x time (within) | Interaction | F (1, 6) = 0.36 | P=0.573 |
| Time | Main effect | F (1, 6) = 0.55 | P=0.488 |
| Sex: male, n = 7; female, n = 5 | Main effect | F (1, 6) = 2.79 | P=0.146 |
| **B** | | | |
| **AUC for coc (1mg/kg/ml) infusions from first to last** | Two-way RM ANOVA | | |
| Sex (between) x time (within) | Interaction | F (1, 7) = 3.79 | P=0.093 |
| Time | Main effect | F (1, 7) = 1.79 | P=0.223 |
| Sex: male, n =6 ; female, n = 5 | Main effect | F (1, 7) = 2.39 | P=0.166 |
| **C** | | | |
| **AUC for coc (3mg/kg/ml) infusions from first to last** | Two-way RM ANOVA | | |
| Sex (between) x time (within) | Interaction | F (1, 12) = 0.25 | P=0.624 |
| Time | Main effect | F (1, 12) = 0.60 | P=0.453 |
| Sex: male, n = 8; female, n = 6 | Main effect | F (1, 12) = 2.16 | P=0.168 |

**Table S12**. Detailed statistics for Figure S2. Significant p-values are bolded.

| *Comparison* | *Statistical Test* | *Statistic* | *p-value* |
|---|---|---|---|
| **B** | | | |
| Maximum Area Under the Curve for Repeated Cocaine 1mg/kg | Two-way RM ANOVA | | |
| Repeated cocaine (within) x Sex (between) | Interaction | F (1, 22) = 0.37 | P=0.547 |
| Repeated cocaine | Main effect | F (1, 22) = 4.72 | **P=0.041** |
| Sex: male, n = 13; female, n = 12 | Main effect | F (1, 22) = 2.03 | P=0.169 |
| **C-D** | | | |
| Area Under the Curve for Repeated Cocaine 1mg/kg | Three-way RM ANOVA | | |
| Time (within) x Repeated cocaine (within) x Sex (between) | Interaction | F (2, 44) = 4.15 | **P=0.022** |
| Time x Sex | Interaction | F (2, 44) = 0.67 | P=0.519 |
| Time x Repeated cocaine | Interaction | F (2, 44) = 35.99 | **P<0.001** |
| Sex x Repeated cocaine | Interaction | F (1, 22) = 1.16 | P=0.293 |
| Time | Main effect | F (2, 44) = 7.74 | **P=0.001** |
| Sex: male, n = 13; female, n = 12 | Main effect | F (1, 22) = 0.36 | P=0.554 |
| Repeated cocaine | Main effect | F (1, 22) = 0.00 | P=0.979 |
| **C** | | | |
| Male Area Under the Curve for Repeated Cocaine 1mg/kg | Two-way RM ANOVA | | |
| Time x Repeated cocaine: n = 13 | Interaction | F (2, 22) = 24.48 | **P<0.001** |
| Time | Main effect | F (2, 22) = 4.28 | **P=0.027** |
| Repeated cocaine | Main effect | F (1, 11) = 0.48 | P=0.502 |
| Repeated cocaine: 0-30s | Holm-Šídák | | **P=0.048** |
| Repeated cocaine: 30-60s | Holm-Šídák | | P=0.502 |
| Repeated cocaine: 60-90s | Holm-Šídák | | **P<0.001** |
| **D** | | | |
| Female Area Under the Curve for Repeated Cocaine 1mg/kg | Two-way RM ANOVA | | |
| Time x Repeated cocaine: n = 12 | Interaction | F (2, 22) = 11.53 | **P<0.001** |
| Time | Main effect | F (2, 22) = 3.99 | **P=0.033** |
| Repeated cocaine | Main effect | F (1, 11) = 0.72 | P=0.415 |
| Repeated cocaine: 0-30s | Holm-Šídák | | **P<0.001** |
| Repeated cocaine: 30-60s | Holm-Šídák | | P=0.967 |
| Repeated cocaine: 60-90s | Holm-Šídák | | P=0.206 |
| **F** | | | |
| Maximum Area Under the Curve for Repeated Cocaine 3mg/kg | Two-way RM ANOVA | | |
| Repeated cocaine (within) x Sex (between) | Interaction | F (1, 20) = 1.17 | P=0.292 |
| Repeated cocaine | Main effect | F (1, 20) = 5.09 | **P=0.036** |
| Sex: male, n = 12; female, n = 10 | Main effect | F (1, 20) = 0.00 | P=0.987 |
| **G-H** | | | |
| Area Under the Curve for Repeated Cocaine 3mg/kg | Three-way RM ANOVA | | |
| Time (within) x Repeated cocaine (within) x Sex (between) | Interaction | F (2, 40) = 4.66 | **P=0.015** |
| Time x Sex | Interaction | F (2, 40) = 0.055 | P=0.947 |
| Time x Repeated cocaine | Interaction | F (2, 40) = 7.87 | **P=0.001** |
| Sex x Repeated cocaine | Interaction | F (1, 20) = 0.71 | P=0.410 |
| Time | Main effect | F (2, 40) = 12.91 | **P<0.001** |
| Sex: male, n = 12; female, n = 10 | Main effect | F (1, 20) = 0.09 | P=0.762 |
| Repeated cocaine | Main effect | F (1, 20) = 11.08 | **P=0.003** |
| **G** | | | |
| Male Area Under the Curve for Repeated Cocaine 3mg/kg | Two-way RM ANOVA | | |
| Time x Repeated cocaine: n = 12 | Interaction | F (2, 22) = 11.68 | **P<0.001** |
| Time | Main effect | F (2, 22) = 4.66 | **P=0.021** |
| Repeated cocaine | Main effect | F (1, 11) = 9.66 | **P=0.010** |
| Repeated cocaine: 0-30s | Holm-Šídák | | P=0.058 |
| Repeated cocaine: 30-60s | Holm-Šídák | | **P=0.029** |
| Repeated cocaine: 60-90s | Holm-Šídák | | **P=0.010** |
| **H** | | | |
| Female Area Under the Curve for Repeated Cocaine 3mg/kg | Two-way RM ANOVA | | |
| Time x Repeated cocaine: n = 10 | Interaction | F (2, 18) = 2.26 | P=0.133 |
| Time | Main effect | F (2, 18) = 12.37 | **P<0.001** |
| Repeated cocaine | Main effect | F (1, 9) = 2.80 | P=0.129 |
| Repeated cocaine: 0-30s | Holm-Šídák | | P=0.295 |
| Repeated cocaine: 30-60s | Holm-Šídák | | P=0.284 |
| Repeated cocaine: 60-90s | Holm-Šídák | | P=0.295 |

**Table S13**. Detailed statistics for Figure S3. Significant p-values are bolded.

| *Comparison* | *Statistical Test* | *Statistic* | *p-value* |
|---|---|---|---|
| **% Magnitude Change for Male Cocaine (1mg/kg/ml) + Rimonabant (1mg/kg)** | Two-way RM ANOVA | | |
| Time (within) x treatment (between): male repeat cocaine, n =13; male Rimonabant, n = 9 | Interaction | F (2, 40) = 33.96 | **P<0.001** |
| Time | Main effect | F (2, 40) = 26.65 | **P<0.001** |
| Treatment | Main effect | F (1, 20) = 2.48 | P=0.131 |
| Repeated Coc1 vs Repeated Coc1 + Rimon 1: 0-30s | Holm-Šídák | | **P<0.001** |
| Repeated Coc1 vs Repeated Coc1 + Rimon 1: 30-60s | Holm-Šídák | | P=0.789 |
| Repeated Coc1 vs Repeated Coc1 + Rimon 1: 60-90s | Holm-Šídák | | P=0.160 |
| **% Magnitude Change for Female Cocaine (1mg/kg/ml) + Rimonabant (1mg/kg)** | Two-way RM ANOVA | | |
| Time (within) x treatment (between): female repeat cocaine, n =12; female Rimonabant, n = 9 | Interaction | F (2, 38) = 8.18 | **P=0.001** |
| Time | Main effect | F (2, 38) = 5.61 | **P=0.007** |
| Treatment | Main effect | F (1, 19) = 1.31 | P=0.267 |
| Repeated Coc1 vs Repeated Coc1 + Rimon 1: 0-30s | Holm-Šídák | | **P=0.005** |
| Repeated Coc1 vs Repeated Coc1 + Rimon 1: 30-60s | Holm-Šídák | | P=0.915 |
| Repeated Coc1 vs Repeated Coc1 + Rimon 1: 60-90s | Holm-Šídák | | P=0.915 |
| **% Magnitude Change for Male Cocaine (1mg/kg/ml) + Rimonabant (3mg/kg)** | Two-way RM ANOVA | | |
| Time (within) x treatment (between): male repeat cocaine, n =13; male Rimonabant, n = 9 | Interaction | F (2, 40) = 13.58 | **P<0.001** |
| Time | Main effect | F (2, 40) = 43.59 | **P<0.001** |
| Treatment | Main effect | F (1, 20) = 1.760 | P=0.200 |
| Repeated Coc1 vs Repeated Coc1 + Rimon 3: 0-30s | Holm-Šídák | | **P=0.003** |
| Repeated Coc1 vs Repeated Coc1 + Rimon 3: 30-60s | Holm-Šídák | | P=0.838 |
| Repeated Coc1 vs Repeated Coc1 + Rimon 3: 60-90s | Holm-Šídák | | P=0.838 |
| **% Magnitude Change for Female Cocaine (1mg/kg/ml) + Rimonabant (3mg/kg)** | Two-way RM ANOVA | | |
| Time (within) x treatment (between): female repeat cocaine, n =12; female Rimonabant, n = 12 | Interaction | F (2, 44) = 5.25 | **P=0.009** |
| Time | Main effect | F (2, 44) = 25.81 | **P<0.001** |
| Treatment | Main effect | F (1, 22) = 9.11 | **P=0.006** |
| Repeated Coc1 vs Repeated Coc1 + Rimon 3: 0-30s | Holm-Šídák | | **P<0.001** |
| Repeated Coc1 vs Repeated Coc1 + Rimon 3: 30-60s | Holm-Šídák | | **P=0.078** |
| Repeated Coc1 vs Repeated Coc1 + Rimon 3: 60-90s | Holm-Šídák | | **P=0.140** |
| **% Magnitude Change for Male Cocaine (1mg/kg/ml) + MJN-110 (5mg/kg)** | Two-way RM ANOVA | | |
| Time (within) x treatment (between): male repeat cocaine, n =13; male MJN-110, n = 8 | Interaction | F (2, 38) = 18.45 | **P<0.001** |
| Time | Main effect | F (2, 38) = 10.13 | **P<0.001** |
| Treatment | Main effect | F (1, 19) = 4.00 | P=0.060 |
| Repeated Coc1 vs Repeated Coc1 + MJN: 0-30s | Holm-Šídák | | P=0.290 |
| Repeated Coc1 vs Repeated Coc1 + MJN: 30-60s | Holm-Šídák | | **P=0.001** |
| Repeated Coc1 vs Repeated Coc1 + MJN: 60-90s | Holm-Šídák | | **P=0.003** |
| **% Magnitude Change for Female Cocaine (1mg/kg/ml) + MJN-110 (5mg/kg)** | Two-way RM ANOVA | | |
| Time (within) x treatment (between): female repeat cocaine, n =12; female MJN-110, n = 9 | Interaction | F (2, 38) = 12.98 | **P<0.001** |
| Time | Main effect | F (2, 38) = 1.12 | P=0.336 |
| Treatment | Main effect | F (1, 19) = 8.27 | **P=0.010** |
| Repeated Coc1 vs Repeated Coc1 + MJN: 0-30s | Holm-Šídák | | P=0.718 |
| Repeated Coc1 vs Repeated Coc1 + MJN: 30-60s | Holm-Šídák | | **P=0.015** |
| Repeated Coc1 vs Repeated Coc1 + MJN: 60-90s | Holm-Šídák | | **P=<0.001** |
| *Comparison* | *Statistical Test* | *Statistic* | *p-value* |
| **% Magnitude Change for Male Cocaine (3mg/kg/ml) + Rimonabant (1mg/kg)** | Two-way RM ANOVA | | |
| Time (within) x treatment (between): male repeat cocaine, n =12; male Rimonabant, n = 11 | Interaction | F (2, 42) = 0.20 | P=0.817 |
| Time | Main effect | F (2, 42) = 3.73 | **P=0.032** |
| Treatment | Main effect | F (1, 21) = 3.96 | P=0.060 |
| **% Magnitude Change for Female Cocaine (3mg/kg/ml) + Rimonabant (1mg/kg)** | Two-way RM ANOVA | | |
| Time (within) x treatment (between): female repeat cocaine, n =10; female Rimonabant, n = 12 | Interaction | F (2, 40) = 0.21 | P=0.808 |
| Time | Main effect | F (2, 40) = 2.15 | P=0.130 |
| Treatment | Main effect | F (1, 20) = 6.38 | **P=0.020** |
| **% Magnitude Change for Male Cocaine (3mg/kg/ml) + Rimonabant (3mg/kg)** | Two-way RM ANOVA | | |
| Time (within) x treatment (between): male repeat cocaine, n =12; male Rimonabant, n = 7 | Interaction | F (2, 34) = 3.17 | P=0.055 |
| Time | Main effect | F (2, 34) = 0.58 | P=0.567 |
| Treatment | Main effect | F (1, 17) = 2.18 | P=0.158 |
| **% Magnitude Change for Female Cocaine (3mg/kg/ml) + Rimonabant (3mg/kg)** | Two-way RM ANOVA | | |
| Time (within) x treatment (between): female repeat cocaine, n =10; female Rimonabant, n = 7 | Interaction | F (2, 30) = 2.13 | P=0.137 |
| Time | Main effect | F (2, 30) = 0.01 | P=0.986 |
| Treatment | Main effect | F (1, 15) = 8.18 | **P=0.012** |
| **% Magnitude Change for Male Cocaine (3mg/kg/ml) + MJN-110 (5mg/kg)** | Two-way RM ANOVA | | |
| Time (within) x treatment (between): male repeat cocaine, n =12; male MJN-110, n = 10 | Interaction | F (2, 40) = 2.05 | P=0.142 |
| Time | Main effect | F (2, 40) = 0.63 | P=0.539 |
| Treatment | Main effect | F (1, 20) = 1.02 | P=0.324 |
| **% Magnitude Change for Female Cocaine (3mg/kg/ml) + MJN-110 (5mg/kg)** | Two-way RM ANOVA | | |
| Time (within) x treatment (between): female repeat cocaine, n =10; female MJN-110, n = 8 | Interaction | F (2, 32) = 0.49 | P=0.615 |
| Time | Main effect | F (2, 32) = 0.07 | P=0.933 |
| Treatment | Main effect | F (1, 16) = 4.20 | P=0.057 |

**Table S14.** Detailed statistics of two-way RM ANOVAs conducted for individual sexes for magnitude change comparison between repeated cocaine infusions versus repeated cocaine infusions with a cannabinoid-targeting drug. Significant p-values are bolded.
